# Modeling the immunological pre-adaptation of HIV-1

**DOI:** 10.1101/2020.01.08.897983

**Authors:** Christiaan H. van Dorp, Michiel van Boven, Rob J. de Boer

## Abstract

It is becoming increasingly evident that the evolution of HIV-1 is to a large extent determined by the immunological background of the host. On the population-level this results in associations between specific human leukocyte antigen (HLA) alleles and polymorphic loci of the virus. Furthermore, some HLA alleles that were previously associated with slow progression to AIDS have been shown to lose their protective effect, because HLA-specific immunological escape variants have spread through the population. This phenomenon is known as immunological pre-adaptation. Apart from adapting to human immune responses, the set-point virus load (SPVL) of HIV-1 is thought to have evolved to values that optimize the population-level fitness of the virus. This suggestion is supported by considerable heritability of the SPVL. Previous modeling studies show that whether or not SPVL optimization is expected to occur depends sensitively on the underlying assumptions with respect to the extent of within-versus between-host selection. Here we use a detailed and semi-realistic multi-level HIV-1 model in which immunological pre-adaptation and SPVL evolution can emerge from the underlying interactions of the virus with the immune system of the host. This enables us to study the effect of immunological escape on disease progression, and how disease progression may be molded by SPVL evolution. We find that the time to AIDS could decrease significantly (0.5-1.0 years) in a HLA-dependent manner by immunological pre-adaptation over the long-term course of the epidemic (> 100 years). We find that SPVL is not expected to evolve to optimize the population-level fitness of HIV-1, even though high heritability of the SPVL emerges from continual selection of immune-escape mutations.

## Introduction

Over the past decades, the evolution of the virulence of Human immunodeficiency virus type 1 (HIV-1) has been studied extensively. High virulence is related to fast disease progression, but also to high set-point virus load (SPVL) and increased infectiousness. A corollary is a life-history trade-off between infectiousness and the duration of the infection, which has been hypothesized to steer the evolution of virulence and SPVL [29]; a hypothesis for which there is mounting evidence [8, 9]. Complicating this idea is the fact that selection operates at different levels [30, 40], which shapes HIV-1’s fitness landscape in a non-trivial manner. Recently, the characteristics of this fitness landscape itself have become a subject of study, wherein the Ising model is used to map HIV genomes to (proxies of) fitness [5, 27, 41, 48, 52]. In addition, the environment (i.e. the host) impacts the shape of the fitness landscape, as evidenced by strong virus-host interactions [61].

Due to genetic variability of the host, notably HLA-polymorphism, the fitness of a single HIV-1 strain can differ markedly between two hosts. After every transmission event new immune responses are to be escaped, while escapes from the transmitter’s cytotoxic T lymphocytes (CTLs) may be reverted whenever such escape mutations have deleterious fitness effects. Not all escape mutations are reverted after transmission, as is clear from analyses of HLA footprints on viral populations world-wide [45]. In these cases, compensatory mutations may have (partially) restored viral fitness. Further, many of these mutations have epistatic effects, and many epistatic interactions are in fact host-mediated, as evidenced by the observation that the co-occurrence of certain mutations can be associated with the HLA type of the host [12]. This process of escape, reversion and compensation drives the evolution of HIV-1 over a complex and rugged fitness landscape.

What is meant by fitness in earlier studies depends on the context. For instance, the virus replicative capacity (vRC; the rate of exponential growth), established by *in-vitro* experiments, is often used as a within-host fitness measure (the Malthusian fitness). On the population level, the expected number of secondary cases caused by one infected individual is thought to be a good notion of fitness, and is equal to the basic reproduction number (ℛ_0_) at the onset of the epidemic. In each case, fitness is the product of viral and host traits. Non-linear and highly stochastic within-host processes determine how individual-level fitness is related to population-level fitness. Here, we model such processes in detail, to gain insight in how the levels of selection are connected.

A number of multi-level HIV-1 models have been developed before. Notably, Lythgoe *et al*. [57] experiment with rugged fitness landscapes, and a dynamic within-host model. Doekes *et al*. [24] extended this model with a reservoir of latently infected cells. Hool *et al*. [42] use a within-host model of HIV-1 to determine the within-host evolutionary dynamics, and population-level parameters. These studies try to reconcile within-host evolution and selection with the adaptive character of SPVL evolution. Previously, we have modeled HIV-1 virulence evolution in a HLA-polymorphic population [26], and found that the population-level SPVL distribution is only significantly shaped by the effects of transmission-adaptation when escape and reversion rates are unreasonably low. Some modeling choices in the previous study were influenced by computational convenience. Here, we follow a bottom-up approach that addresses the more *ad hoc* simplifications made before. Specifically, the within-host processes are modeled in detail, and the higher level variables are derived from the within-host model.

In order to couple a between-host with a within-host model, the infectiousness and the duration of the infectious period need to be derived from within-host processes. For the infectiousness and duration we use the empirical relation with virus load [8, 29]. Hence, we require a model that yields viral load from the within-host dynamics. Our analyses are based on a widely used system of ordinary differential equations [2, 22], describing the relation between target cells (CD4^+^ cells), HIV-1 infected cells, and effector cells (HIV-1 specific CTLs). Apart from the virus load, this model provides us with rules for the introduction of new mutants and the addition of new immune responses. The model allows for clonal interference and slow rates of immunological escape. Both phenomena have been described for CTL escapes [70, 75].

Separately, the key ingredients of our HIV-1 immuno-epidemiological model can be found in the three examples of nested models mentioned before [26, 42, 57]. The model presented here combines all of these components: (i) The parameters determining the viral dynamics are determined by a complex and rugged genotype-to-phenotype mapping, as was touched upon by Lythgoe *et al*. [57]. We improve upon this by using the *NK* model [44] that has been developed as an abstract fitness model with epistatic interactions, and by allowing multiple parameters to be determined by the virus, instead of just the replicative capacity. (ii) The virus load, the resulting epidemiological parameters, and individual-level growth rates are calculated using a realistic model of within-host HIV-1 dynamics, akin to the work of Hool *et al*. [42], which is improved upon by allowing for co-existence of, and competition between multiple viral strains. (iii) Individual-level evolution is directed, but because immune responses are restricted to polymorphic HLA alleles, the effects of a particular mutation can be different in each host [26]. Here, we further add the possibility of compensatory mutations (through the *NK* model), HLA alleles with realistic population-level frequencies, and immuno-dominant and sub-dominant immune responses. Notice that in the case of hepatitis C virus, evolving interactions between a virus and the immune system have been incorporated in a similar multi-level model before [53].

Interest in population-level adaptation of HIV-1 is certainly not restricted to the SPVL. It has been known for a long time that certain HLA molecules are associated with a low SPVL and slow disease progression [46, 50]. For instance the HLA-B*57 and HLA-B*27 types are over-represented in so-called elite controllers: patients with a nearly undetectable virus load in the absence of antiretroviral therapy. In more recent years, advanced statistical methods have been developed to discover associations between HLA molecules and viral polymorphisms [14], revealing widespread footprints of HLA in the genome of HIV-1.

A consequence of widespread escape from HLA-restricted immune responses is that individuals can be infected with viruses that are already adapted to their immune responses, because the same HLA molecule was encountered in a previous host. This phenomenon is referred to as “pre-adaptation” [13]. Pre-adaptation can lead to the apparent loss of HLA associations, if the statistical model does not properly correct for this pre-adaptation [13]. Furthermore, evidence is accumulating that HLA pre-adaptation is a major determinant of disease progression [63], although currently the population-level health effects appear to be relatively small [47].

The spread of CTL escape mutations through the human population has been modeled before, but these models are restricted to only a few epitopes and HLA alleles [31, 32]. In the model presented here, we are not restricted to merely a couple of host and viral variants, and can follow escape mutations on a much larger scale. Furthermore, our bottom-up modeling approach allows us to investigate SPVL evolution, heritability, and the detrimental effects of pre-adaptation in an integrated manner.

## Results

### A multi-level HIV-1 model

Following previous modeling studies [38, 39], we simulate a population of individuals that can infect each other via a dynamic sexual contact network (Figure S1). The rate at which a susceptible individual becomes infected depends on the number of infected partners, and each partner’s transmission rate. The transmission rate of an infected individual, in turn, depends on the current virus load (VL), which is determined by the individual-level model (see below). The relation between VL and transmission rate has been estimated in a cohort of transmission pairs from Zambia [29]. Furthermore, the VL relates to the risk of progression to AIDS, for which we use estimates from a cohort from Amsterdam [29]. In contrary to previous work [26, 29, 77], the acute and AIDS stages of the HIV-1 infection are not explicitly modeled here. Instead, the increased transmission rate during the acute infection naturally results from the VL in our individual-level model, and we remove infected individuals from the population at the end of the asymptomatic phase (i.e. the onset of AIDS).

At the individual level, we model resting and susceptible populations of CD4^+^ T cells (quiescent and target cells), infected cells, free virus, and CD8^+^ T cells (effector cells). We do not explicitly model the B-cell response, and as we are interested in an epidemic of untreated infections, we ignore latently infected cells. The dynamics of these within-host populations is given by a system of ordinary differential equations (ODEs; Equation (1) in Methods). New immune responses and mutant virus strains are added to the system at random points in time, determined by an antigen-dependent immune-activation rate, and the mutation rate of the virus, respectively. An example of an individual-level simulation is given in Figure 1.

**Figure 1:**
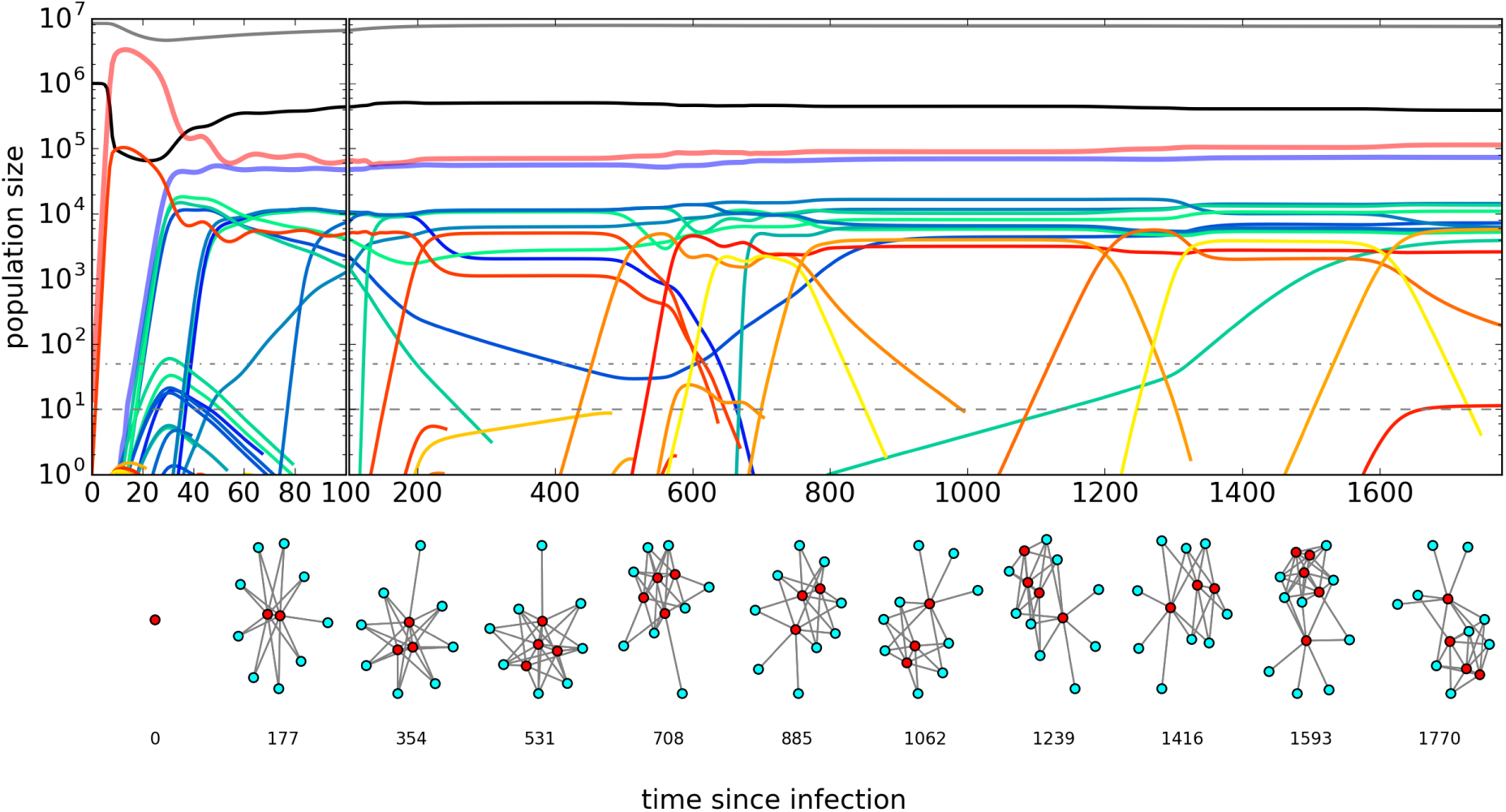
An example of an individual-level simulation. The top panel shows the time series of the population sizes of quiescent CD4^+^T cells (gray), target cells (black), infected cells (yellow-red), and effector cells (green-blue). The thick lines indicate the total virus load (red) and the total number of effector cells (blue). The horizontal dashed line marks the population size at which declining populations are considered endangered (*τ*_end_, see Methods), and the dot-dashed line pinpoints the population size at which we start tracking the mutants of a viral strain (*τ*_mut_, see Methods). The networks indicate the interactions between immune responses (blue) and viruses (red). An immune response is connected with a virus if the virus contains the response’s epitope. Two different viruses are connected if they differ at precisely one locus. Figure S3 shows the individual-level dynamics for different mutation rates, but otherwise identical initial conditions and parameters.

In previous modeling studies, the within-host fitness of the virus is typically modulated by only one or two parameters of the model [e.g. 22, 54]. Large variation in the SPVL can be achieved by allowing many parameters to vary by a small amount [65]. Here, we generalize this idea by putting multiple parameters of the ODE model under control of the virus. The genome of a virus is modeled using a sequence of *N* = 540 bits, and consists of 9 genes of equal length. The genes are responsible for specific functions of the virus. This is accomplished by letting each gene encode the value of a parameter in the system of ODEs. The default parameter values are listed in Table 1; the genes determine the deviation from the default (see Methods).

**Table 1:** Parameters of the individual-level model. Parameters are determined by the host (h), the virus (v) or the immune response (r). This is indicated with an “x” in the respective columns. The given parameter value is either the default value, or a range of possible values. Notes: (1) Given *d*_Q_ and *ω*_0_, we choose *s* such that *Q* = 10^6^mL^−1^ in the average healthy individual. *d*_Q_, *ω*_0_ and *ω*_*v*_ are taken from Bartha *et al*. [4]. (2) The activated CD4 T cells are assumed to be much shorter-lived than resting CD4 T cells. (3) An intermediate virus load to maximize the role of target-cell activation. (4) The parameter *β* is chosen such that the vRC is 1.5 per day [33]. (5) Conservatively taken from Doitsh *et al*. [25]. (6) Taken from Dixit *et al*. [23]. (7) Chosen such that the death rate of an infected cell is 1d^−1^ [59]. (8) The viral clearance rate is taken from Ramratnam *et al*. [72], then we choose a production rate to obtain intermediate virus loads. (9) Given a death rate of 0.1d^−1^, the maximum production rate *p*_E_ is taken such that the growth rate of *E* is 1d^−1^ [21]. (10) The parameter *λ*_naive_ is chosen such that viable responses are activated subsequently during the acute phase of the infection, and λ_memo_ is chosen such that viable memory is activated within a few days. (11) By taking 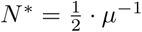, the point-mutation rate (*µ*_*v*_*q*_*v*_) is maximized at 2*µ*. Therefore, a mutation rate *µ*_*v*_ larger than 2*µ* does not increase the point-mutation rate.

| symbol | description | h | v | r | value | unit | note |
| --- | --- | --- | --- | --- | --- | --- | --- |
| $s$ | production of quiescent target cells | x | . | . | $1.4 \cdot 10^4$ | cells mL<sup>-1</sup>d<sup>-1</sup> | (1) |
| $d_Q$ | death rate of uninfected quiescent target cells | x | . | . | $10^{-3}$ | d<sup>-1</sup> | (1) |
| $d_T$ | death rate of uninfected activated target cells | x | . | . | 0.1 | d<sup>-1</sup> | (2) |
| $\omega_0$ | activation rate of quiescent cells | x | . | . | $1.2 \cdot 10^{-2}$ | d<sup>-1</sup> | (1) |
| $\omega_v$ | virus-induced activation rate of quiescent cells | x | x | . | $1.2 \cdot 10^{-2}$ | d<sup>-1</sup> | (1) |
| $h_Q$ | Michaelis-Menten parameter of quiescent cells | x | . | . | $3.3 \cdot 10^4$ | virions mL<sup>-1</sup> | (3) |
| $\beta_v$ | infection rate of target cells | x | x | . | $1.37 \cdot 10^{-6}$ | virions<sup>-1</sup> mL d<sup>-1</sup> | (4) |
| $f_v$ | fraction of non-abortive target cell infections | x | x | . | 0.1 | - | (5) |
| $\mu_v$ | mutation probability of a single locus | x | x | . | $10^{-6}$ | - | - |
| $\gamma_v$ | eclipse-to-production rate | x | x | . | 1.0 | d<sup>-1</sup> | (6) |
| $\alpha_v$ | virus-induced cytopathic effect | x | x | . | 0.9 | d<sup>-1</sup> | (7) |
| $m_v$ | virus-regulated MHC expression | . | x | . | 1.0 | - | - |
| $k_r$ | killing rate of virus-producing cells | . | . | x | $10^{-5} - 10^{-4}$ | cells<sup>-1</sup> mL d<sup>-1</sup> | - |
| $p_{V,v}$ | production rate of virions | x | x | . | 1150 | virions d<sup>-1</sup> | (8) |
| $d_{V,v}$ | clearance rate of virions | x | x | . | 23.0 | d<sup>-1</sup> | (8) |
| $p_E$ | maximal production rate of effector cells | x | . | . | 1.1 | d<sup>-1</sup> | (9) |
| $h_{E,r}$ | Michaelis-Menten parameter of effector cells | . | . | x | $10^1 - 10^5$ | cells mL<sup>-1</sup> | - |
| $d_E$ | death rate of effector cells | x | . | . | 0.1 | d<sup>-1</sup> | (9) |
| $\lambda_{\text{naive}}$ | activation rate of naive CD8<sup>+</sup> T cells | x | . | . | 0.1 | d<sup>-1</sup> | (10) |
| $\lambda_{\text{memo}}$ | activation rate of memory CD8<sup>+</sup> T cells | x | . | . | 1.0 | d<sup>-1</sup> | (10) |
| *compound parameters* |  |  |  |  |  |  |  |
| $q_v$ | probability of faithful reverse transcription | x | x | . | $e^{-N^*\mu}$ | - | (11) |

The most obvious parameter under control of the virus is the infection rate of target cells (*β*). The gene encoding *β* is analogous to the *env* gene that encodes the gp120 and gp41 proteins that mediate the attachment and entry to the target cell. After entry of the target cell, the infected cell is in an eclipse phase before it starts producing virions. The length of the eclipse phase (regulated by the parameter *γ*) is determined by another gene. This gene is analogous to the *pol* gene that encodes reverse transcriptase (RT) and integrase. The integration of the viral genome into the host cell’s DNA has a high probability of failing, which is referred to as abortive infection, often leading to apoptosis or pyroptosis of the cell [20]. We model this with a virus-controlled parameter *f*, representing the fraction of successfully infected cells. The gene encoding *f* is partly analogous to *vif*, which disrupts the innate intra-cellular APOBEC3 response [73].

Next, we have parameters for production (*p*_V_) and degradation (*d*_V_) of virions. Although many genes are involved in the formation of virions, one can think of the *tat* gene that regulates transcription. Degradation of virions is modulated by the stability of structural proteins, for instance p17 and p24, encoded by the *gag* gene. Variation in *d*_*V*_ can further be caused by differences in antibody affinities and concentrations. Even in the absence of an immune response, an infected cell experiences a virus-induced cytopathic effect [20]. We model this using a virus-controlled parameter *α*, which is added to the death rate of virion producing cells. The gene encoding *α* could be analogous to the *nef* gene. The same *nef* gene accounts for the down-regulation of MHC. We model the expression of MHC using a virus-controlled parameter *m*, which is used to regulate the killing rate of the infected cell by effector cells.

HIV-1 is known to modulate the activation of CD4^+^ T-cells, making them permissive to infection. This principle has been used before in modeling studies [4, 42], and in our model we use a virus-controlled parameter *ω* to boost the activation of quiescent CD4^+^ T-cells. It is not well known how exactly HIV-1 influences the activation of CD4^+^ T cells, and although many candidate mechanisms exists [4], in our model, the parameter *ω* is determined by a single gene.

Finally, the mutation rate (*µ*) is modulated by the virus in analogy with the RT protein. The process of reverse transcription is highly error-prone and results on average in one mutated nucleotide per infected cell [58]. It has even been suggested that HIV-1 and other RNA viruses evolve close to the error threshold [80], which means that an increase in *µ* should result in a significant fitness cost. However, to make the simulations computationally feasible, it is impossible to simply choose *µ* = 1*/N* as a default value of the mutation rate and we are forced to take a much smaller default value (e.g. *µ* = 10^−6^). To simulate the nearby error threshold, and prevent viruses from evolving very large mutation rates, we let *N* * *> N* denote the length of the full genome, and think of the *N* modeled loci as a small subset of all the *N* * loci. Mutations in the *N* * − *N* un-modeled loci are assumed to be highly deleterious, which we implement by multiplying the infection rate *β* with the probability of faithful replication 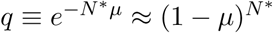 in Equation (1).

The relation between genotype and parameter value (phenotype) is given by the *NK* model [44], which generalizes the Ising model that has been used previously to infer properties of the fitness landscape of HIV-1 [5, 27, 41, 48, 52]. The *NK* model has been applied in many different settings, including immunological models [18], and allows for epistatic interactions between loci, which means that the effect of a mutation depends on the genetic background in which the mutation occurs. The exact implementation of the *NK* model is given in the Methods section. In addition to the parameter variation determined by the virus, each host has slightly different (10% standard deviation) individual-level parameters, to account for host-effects that are not due to the HLA genotype.

The virus- and host-dependent parameters each have a different effect on the SPVL. Figure 2 shows a cross-sectional sample from a simulated epidemic (see below). The SPVL is defined as the average virus load over the infectious period (see Figure 2A for individual time series). The gene with the largest effect on the SPVL modulates the *β* parameter, but also the genes responsible for MHC down-regulation (*m*), virion production (*p*_*V*_), virion degradation (*d*_*V*_), and abortive infection (*f*) are important factors. The length of the eclipse phase (*γ*), the mutation rate (*µ*) and target cell activation (*ω*) only have a limited effect on the SPVL. From the side of the host, the parameters that influence the SPVL the most are related to the number of target cells: the influx of CD4^+^ T-cell (*s*) and the death rate of uninfected target cells (*d*_T_), which is in agreement with previous model predictions [10]. Other important host-dependent parameters are related to the immune response [68]. Indeed, the maximum proliferation rate (*p*_E_) and the death rate (*d*_E_) of effector cells decrease and increase the SPVL, respectively.

**Figure 2:**
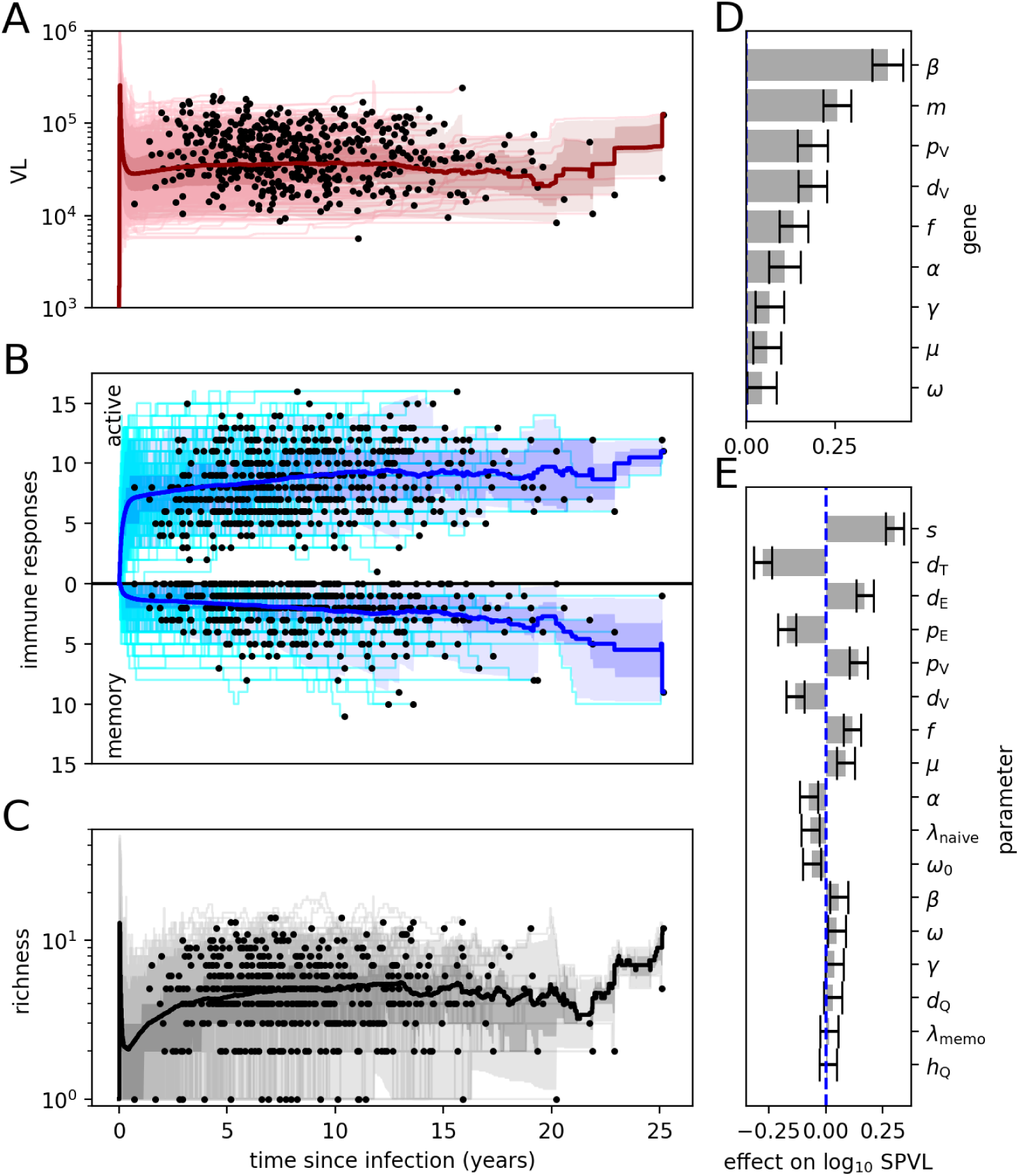
Individual-level time series from a cross-sectional sample. The sample (of size 500) is taken 200 years into an epidemic. The colored bands indicate the 2.5, 25, 75, and 97.5 percentiles, the thick line the average. The end of an infection is indicated with a black dot. **A**. Time series of the virus load (VL). **B**. Time series of the number of activated immune responses. The number of responses that are in the memory state (see Figure S2) are mirrored in the horizontal axis. **C**. Time series of the individual-level diversity, showing the number of strains (richness). **D**. The effect of the genes on the SPVL. A gene encodes an auxiliary parameter (*w*) using the *NK* genotype-to-phenotype map (see Methods). This *w* is used to scale the default value of a particular parameter in the system of ODEs. The effect on the standardized log_10_ SPVL of the *w* parameter of each gene is estimated with multiple linear regression. The error bar indicates the standard error or the estimate. **E**. The effect of the random host effects on the SPVL. The individual-level parameters are slightly different for each host. The effect of this variation on the standardized log_10_ SPVL is again estimated with multiple linear regression.

It is long known that immuno-dominance plays an important role in control of HIV-1 [69]. Furthermore, modeling suggests that the successive emergence of immune responses could aid early escape [22]. We therefore include both phenomena in our model. The system of ODEs specifying our model has the property that escape becomes exceedingly difficult when the immune response is broad, since the contribution of each single response to the total killing rate is small [22, 33]. It follows that equipping each host with too many responses in an early stage of the infection would retard viral escape and therefore individual-level evolution. On the other hand, limiting the number of responses leads to the unnatural situation where during many infections all responses are escaped. This problem is solved with immuno-dominance.

In the individual-level model, a number of CTL responses is in an *active* state, and these responses contribute to the killing rate. The remainder of potential responses can either be *trivial*, i.e. the virus does not (yet) express the epitope, or be in a *latent* state. The latent responses can become activated (at rate *λ*_naive_) when e.g., an active response is escaped (Figure S2), and therefore the number of infected cells increases. When the antigen targeted by an escaped immune response re-appears, the escaped response, that has been maintained in a memory state, can become re-activated (Figure S2). This happens at a faster rate than the activation of naive CD8^+^ T cell (*λ*_memo_ ≫ *λ*_naive_).

In has been shown recently that after escape from one CD8^+^ T-cell response, HIV-1 can induce *de-novo* CD8^+^ T-cell responses [36]. In order to facilitate this behavior, each immune response targets a specific epitope, which is encoded by a locus in the genome and a particular combination of 3 bits. This is a simplification of a peptide epitope, usually 9 amino-acids long, for which the binding affinity to the HLA molecule is mostly determined by two anchor residues, and 6 other residues interact with the complement-determining region of the T-cell receptor (TCR). In our simplification, 1 bit determines HLA-binding, and 2 bits determine TCR-binding. In other words, the first bit determines if an immune response can be mounted against the epitope, and the remaining bits specify the response.

In a similar fashion to viruses determining parameters of the system of ODEs, immune responses control some of these parameters as well. For the rate of expansion of a CTL response, we use the “Beddington” functional response [6, 11] which has recently been shown to explain the power-law dependence of the fold expansion of the CTL response on the precursor number [60]. Immuno-dominance ranking is modeled by varying the amount of antigen needed for immune-response activation (*h*_E_). The second parameter determined by the response is the killing rate (*k*). As we expect that immuno-dominant responses are also efficient at killing infected cells, the parameters *h*_E_ and *k* are negatively correlated.

The number of immune responses that is mounted varies between individuals and over the course of the infection (Figure 2B; active). A part of the activated responses are escaped or out-competed, and are de-activated (Figure 2B; memory). However, not every escape leads to subsequent de-activation of the immune response, as the virus population rapidly becomes diverse (Figure 2C), with each sub-population controlled by a different subset of the active immune responses (Figure 1; networks).

Although the number of target cells is not used to determine disease progression, ongoing escape from CD8^+^ T-cell responses leads to a slow decline in the target cell count (Figure S4), which has been interpreted as the onset of AIDS in similar models [22]. In agreement with data [74], the slope of the CD4^+^ T-cell count (ΔCD4) is negatively correlated with the SPVL (Figure S4B). Interestingly, the relation between ΔCD4 and SPVL appears to be non-linear, which has also been noticed by Regoes *et al*. [74]. A simple explanation for this non-linearity is given by the fact that ΔCD4 is naturally bounded from above: uninfected individuals and elite controllers have a ΔCD4 ≈ 0.

### Immunological escape generates genetic variation and heritability of the SPVL

When HIV-1 is transmitted to a new population, it encounters a set of new HLA molecules. To accommodate for this environmental change, we seed the simulated epidemic with a virus that is pre-evolved in a burn-in simulation. The burn-in simulation has identical parameters, except that the set of HLA molecules is different. The basic population-level statistics of a simulation of the secondary population are shown in Figure 3. The average vRC and SPVL are relatively stable. The SPVL slightly increases, presumably because of immune escape (see below). The SPVL is highly variable although the range is still two orders of magnitude lower than ranges observed in human populations. During the epidemic, the virus drifts away from the initial genotype (wild type), while the genetic variation increases (Figure 3E). The increase in genetic variation coincides with an increase of the heritability of the SPVL, which reaches values around 20% (Figure 3D).

**Figure 3:**
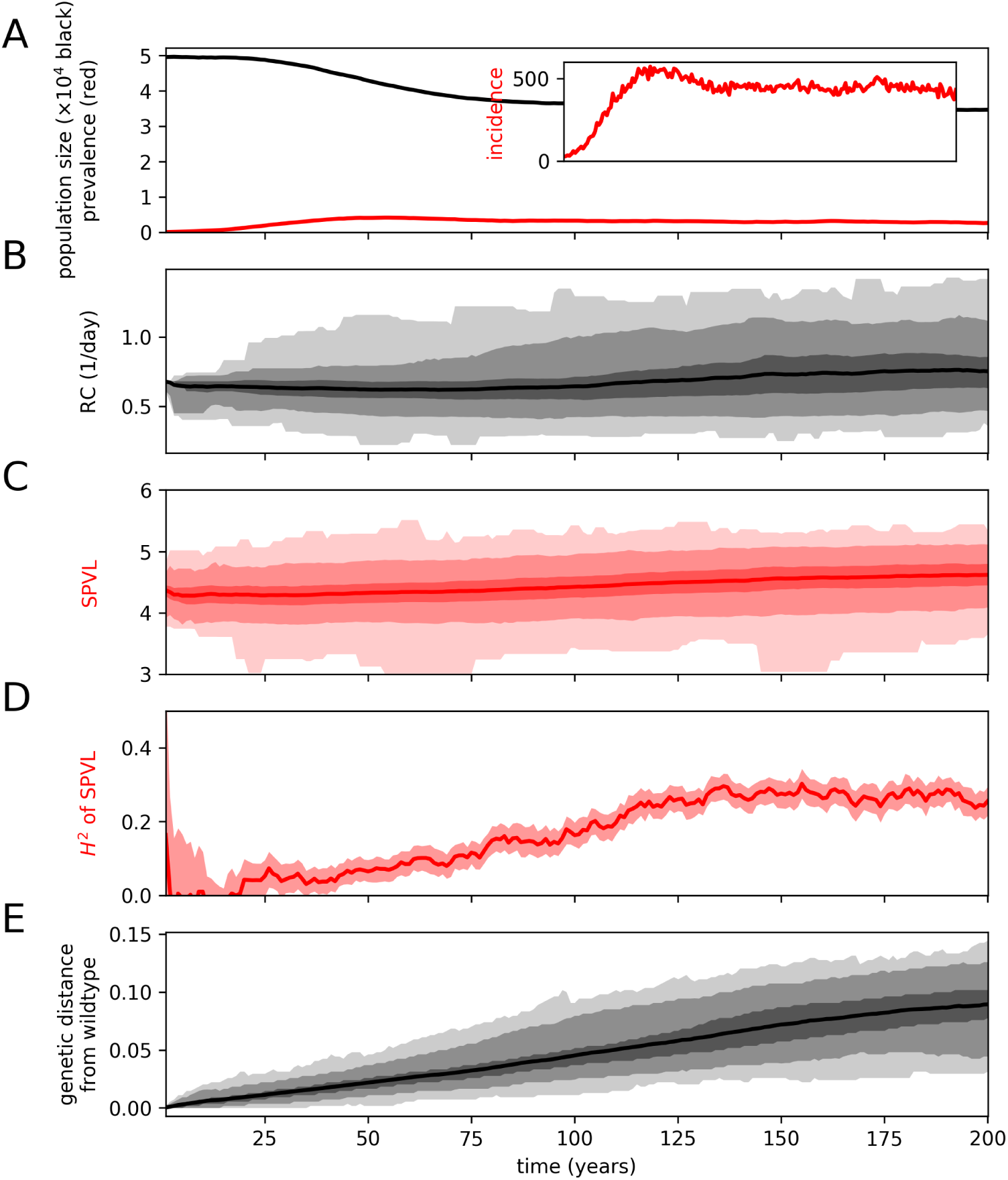
Statistics of the population-level simulation. **A**. Population size (black), prevalence of HIV-1 infected individuals (red) and incidence per year (inset) **B**. The generic replicative capacity (vRC) of the virus at the time of infection **C**. The set-point virus load (SPVL) **D**. The heritability (*H*^2^) of the SPVL. **E**. The genetic (Hamming) distance from the wild-type virus (scaled by genome size). The diversity per locus is presented in Figure S5. The colored bands in panels **B, C** and **E** correspond to the 0, 2.5, 25, 75, 97.5 and 100th percentiles, and the band in panel **D** corresponds to the 95% confidence interval (computed by bootstrapping). The thick lines in panels **B, C, D** and **E** correspond to the mean values.

The relation between SPVL, infectiousness and the duration of the chronic phase results in a life-history trade-off. Hosts with an intermediate SPVL cause the most secondary infections. The fact that SPVL is heritable may result in the optimization of the basic reproduction number (ℛ_0_) by means of SPVL evolution. The first evidence [29] that SPVL has evolved to optimize ℛ_0_ comes from comparing the cross-sectional SPVL distribution with the shape of the transmission potential (TP; i.e. *β*(*V*) × *D*(*V*)). Corroborating evidence comes from a longitudinal study [8], that shows that the mean SPVL changes over time, presumably towards an optimal value. The latter study also suggests that the location of the transmission potential can differ between populations.

In order to test if similar behavior can be observed in our model, we conducted 5 simulations, starting with identical wild-type viruses and populations, but with slightly shifted TPs. The expectation is that the distribution of SPVLs approaches the shape of each TP. However, this expectation is not met in our model (Figure 4A), which is in line with the notion of short-sighted evolution [56, 57] and our previous work [26]. Although optimization of the reproduction number seems intuitively obvious, it is not necessarily the expected behavior from an evolutionary perspective [51, 66]. We therefore extended our *NK* model with a mutation-at-transmission model for the SPVL [39, 77] to verify that under neutral individual-level evolution, ℛ_0_-optimization is indeed expected to occur (see Methods). In the 5 simulations with neutral SPVL evolution, the mean log_10_ SPVL evolves towards the ℛ_0_-optimizing value (Figure 4B), confirming that our population-level model does not preclude ℛ_0_-optimization.

**Figure 4:**
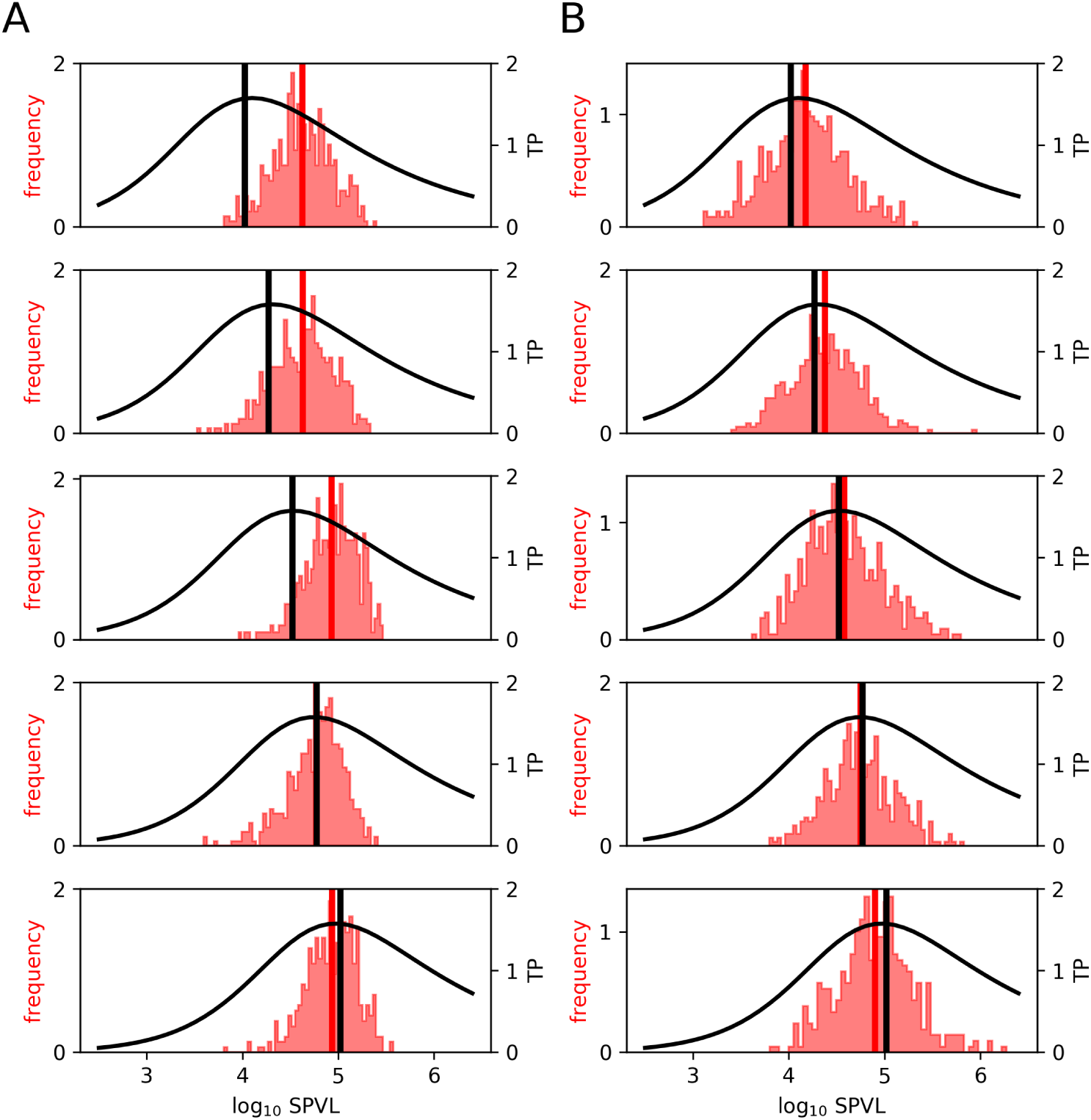
The SPVL distribution is insensitive to the location of the transmission potential. **A**. The 5 panels show the SPVL distribution after 300 years of evolution in identical populations (red histograms), except for the location of the transmission potential (TP; black curves). In each panel, the population-level transmission rate *β* is shifted and scaled, resulting in transmission potentials with optima of identical height. However, these optima are attained at different SPVL values (black vertical lines). The population mean SPVLs are indicated by red vertical lines, **B**. The procedure is identical as in panel **A**, but in this case the mutation-at-transmission model from Shirreff *et al*. [77] was included (see Methods).

Even though ℛ_0_ is not optimized in our model, the virus is clearly subject to genetic drift as the genetic distance form the wild-type virus, as well as the genetic diversity in the population steadily increases as the epidemic progresses (Figure 3E). A closer look at the genetic diversity at each individual locus of the virus shows that polymorphisms preferentially arise at loci that are targeted by frequent and immuno-dominant immune responses (Figure S5D). At the same time, loci at which mutation leads to a high fitness cost (as measured by vRC) are generally more conserved (Figure S5E). Hence, selection pressure from CD8^+^ T-cell responses leads to a genetically diverse virus population at the epidemic level. This genetic variability impacts the individual-level parameters, which results in a positive heritability of the SPVL (Figure 3D).

### Immunological pre-adaptation reshapes the HLA-dependent time to AIDS

HIV-1 polymorphisms are known to be associated with specific HLA alleles, which is a result of the adaptation of HIV-1 to CD8^+^ T-cell responses [64]. These associations are lost over time as escape mutations spread through the population [49]. By correcting for the phylogenetic structure of the viral population, these lost associations can be recovered [12, 14, 31]. In order to see if this form of adaptation occurs in our model, we calculate every 5 years the odds ratios of observing viral loci that are mutated away from the wild-type virus given the presence of a particular HLA allele (see Methods). All significant associations are shown in Figure 5A. Many of these associations are only short lived, and are only observable during the first 50-100 years of the epidemic. However, when we correct for which allele was transmitted to the host (conveniently, we do not have to reconstruct a phylogeny to infer the transmitted allele [12]) many of the short-lived associations become long-lived (Figure 5B). Hence, we find the same statistical clues of pre-adaptation in our simulations that have been observed in real HIV-1 epidemics.

**Figure 5:**
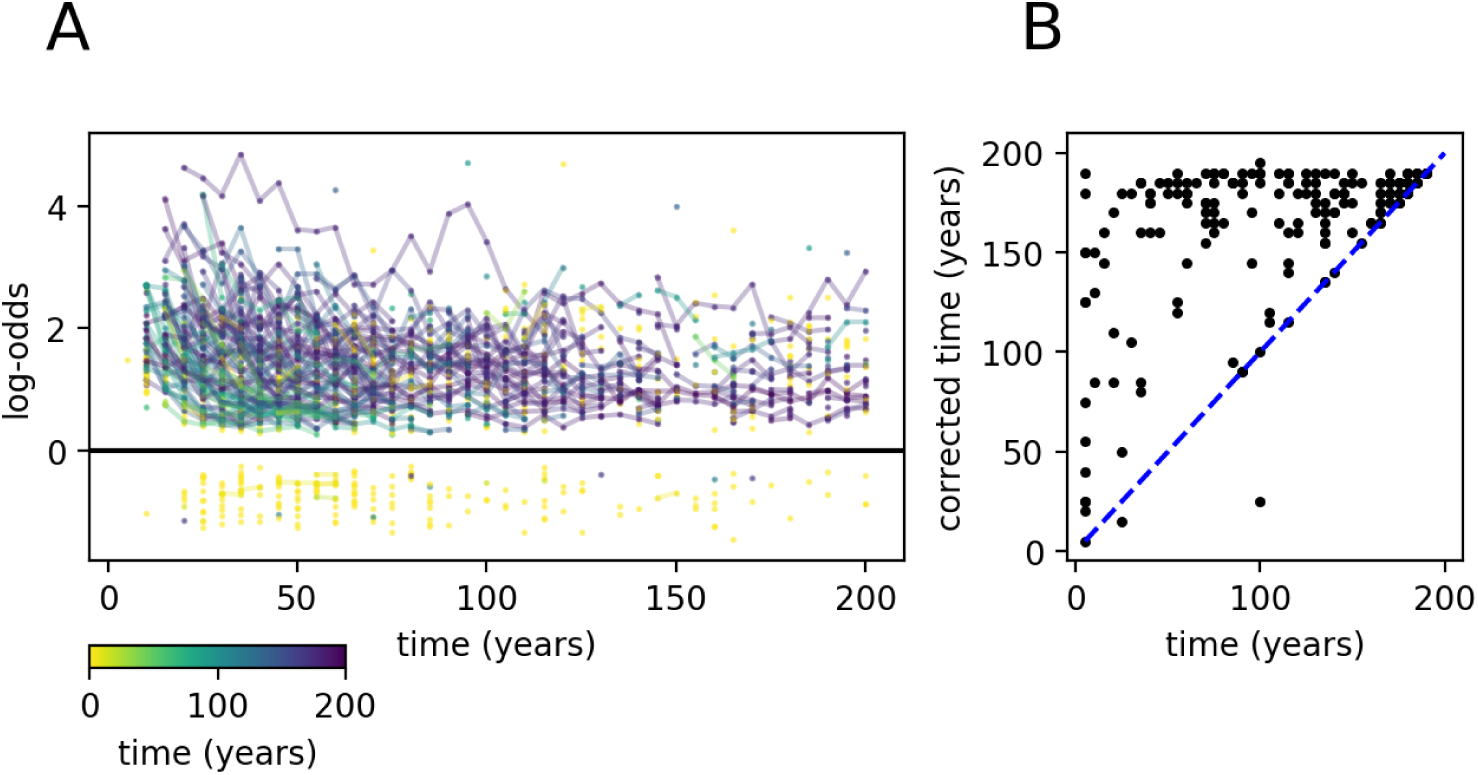
Associations between HLA alleles and viral polymorphisms disappear. **A**. Time series of the strength (log odds ratios) of associations between HLA alleles and viral polymorphisms during a simulated epidemic. The color of each time series corresponds to the length of the time interval in which the association is called significant. **B**. The time that an association remains significant increases after correction for the transmitted virus. The length of the time interval in which an association was called significant is shown without (*x*-axis) and with (*y*-axis) this correction.

The spread of CTL-escape mutations through the population can have a significant impact on virulence, in a way that is dependent on the HLA genotype of the patient [13, 19, 47, 71]. Having full knowledge of the transmitter-founder virus, the exact genetic makeup of the virus that seeded the epidemic (the wild type), and all possible responses that a host is able to mount, it is possible to quantify the impact of pre-adaptation on progression to AIDS in our model. Every 20 years, we make a clone of each infected individual (with identical HLA alleles and host-specific parameters), and infect this clone with the wild-type virus. We define the impact of pre-adaptation as the difference between the lengths of the infections of the clone and the original (Figure 6A; Δ length). To quantify pre-adaptation, we enumerate all possible immune responses against the transmitted virus, and compare this to all possible immune responses the clone can mount to the wild-type virus. The pre-adaptation is defined as the relative difference between the number of possible immune responses (weighted by immuno-dominance). This gives the extent to which the transmitted virus is pre-adapted to the receiving host as compared to the wild type.

**Figure 6:**
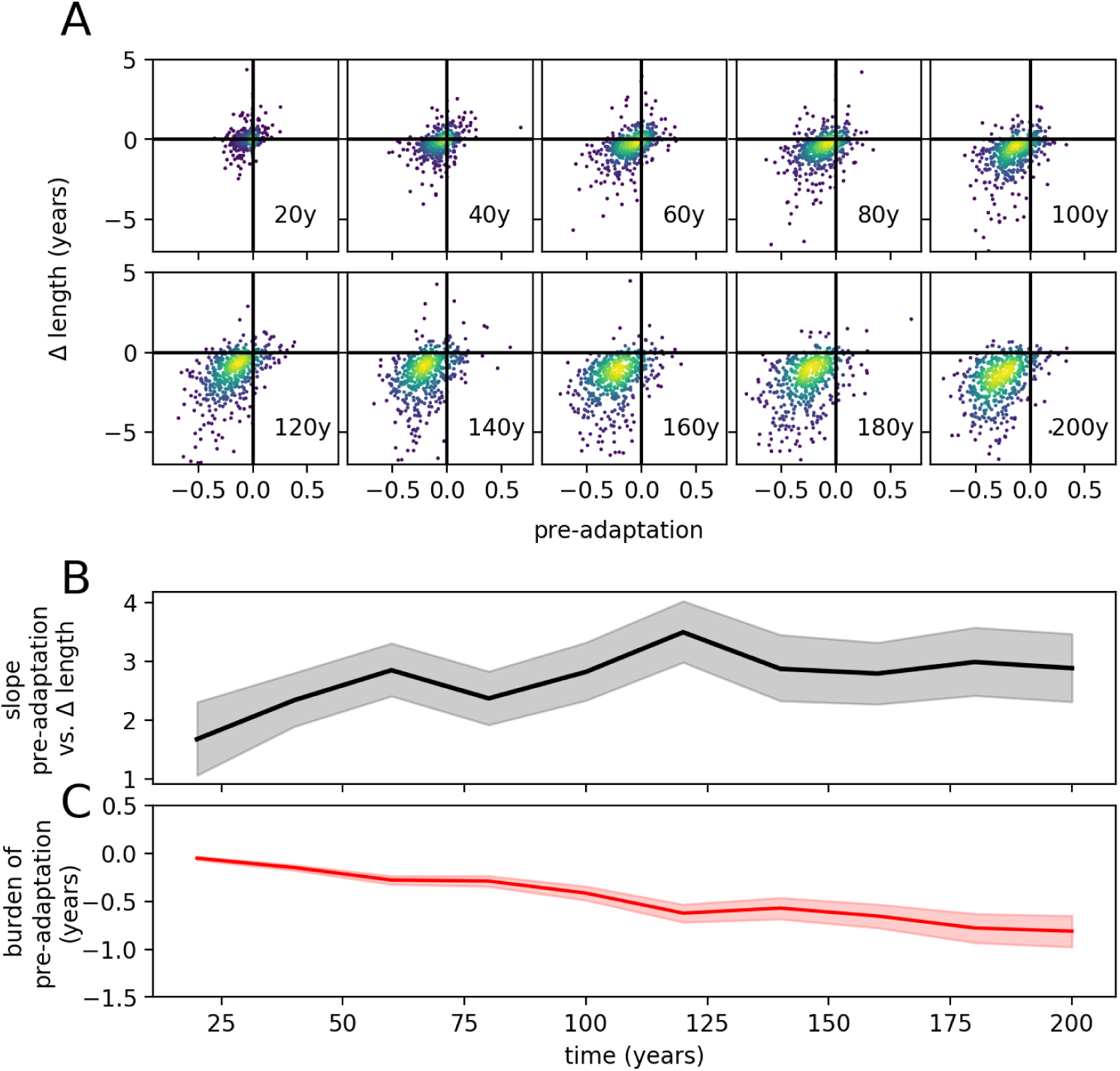
The burden of immunological pre-adaptation increases during the epidemic. **A**. The scatter plots show the relation between relative pre-adaptation of the TF virus with respect to the WT virus, and the difference in the duration of the infection before progression to AIDS. The plots were made at regular time intervals. The color indicates the density of the observations. **B**. At each time point, a slope is calculated with linear regression, which are displayed in the middle panel. The slope indicates the impact of a unit of pre-adaptation in terms of years lost. The band indicates the 95% CI. **C**. The bottom panel shows the average impact of pre-adaptation. This value (and the 95% CI) is calculated by multiplying the average pre-adaptation with the slope of the linear regression.

As the epidemic progresses, the virus escapes from common and immuno-dominant immune responses, which coincides with a faster progression to AIDS (Figure 6A). The relation between pre-adaptation and increase in virulence becomes more pronounced as both become more variable (Figure 6B). The overall effect of pre-adaptation is shown in Figure 6C, and is relatively small: the progression to AIDS is 0.5-1 year faster after 200 years of evolution. This is because most individuals are infected with a virus that is ≈ 30% pre-adapted. These results are roughly comparable with real data: On the one hand, the effect of being infected with a pre-adapted virus can be quite large [63]. However, the fraction of infections with highly pre-adapted viruses currently remains low [47]. A warning from this study would be that the level of pre-adaptation is expected to increase during the next century.

## Discussion

We have developed a multi-level model of HIV-1 that incorporates a number of realistic attributes. The most notable features are: (i) A virus is represented by a genome that encodes multiple parameters of the within-host dynamics. (ii) The individual-level model allows for competition between multiple strains and escape from various immune responses that vary in immuno-dominance. (iii) The host population is highly heterogeneous, but individuals can share immune responses due to pre-defined HLA alleles. As a consequence, the model allowed us to jointly study two population-level evolutionary aspects of HIV-1; i.e. evolution of the SPVL and adaptation to CD8^+^ T-cell responses.

The model behaves realistically when it comes to the spread of escape mutations. Associations between HLA alleles and viral polymorphism arise quickly at the start of the epidemic, and subsequently disappear due to pre-adaptation. Infections with pre-adapted virus result in faster disease progression, and we predict that this impact of pre-adaptation will continue to grow. Previously, we argued that because of short-sighted evolution within the host, SPVL evolution towards ℛ_0_-optimizing values is not to be expected when the mutation rate is high. This was true even in a population with massive HLA polymorphism [26]. However, we left open the possibility that ℛ_0_-optimization could still emerge in a model with a more intricate fitness landscape that allows for within-host competition between strains. Here we have tested this idea, and have to conclude that our original expectation remains valid. These observations potentially have important public-health implications. First, the potential of HIV-1 for pre-adaptation might negatively impact the long-term success of promising HIV-1 vaccines that are currently under development [3]. Second, the risk that HIV-1 virulence increases as antiretroviral treatment becomes more widely available might be limited, contrary to previous model predictions [38, 78].

Although our within-host model is already relatively complex, it still misses a number of important aspects of HIV-1 dynamics. For instance, we do not explicitly model antibody responses, even though the antibody response might be a key factor in the breakdown of control of the virus [81]. Another ignored hallmark of HIV-1 is recombination. Recombination has been included in models that study, e.g., the decrease of the immune-escape rate with the time since infection [34, 35], as a result of clonal interference. Efficient recombination reduces the effect of clonal interference, thereby increasing the rate of individual-level adaptation by means of escape, reversion, and compensation [67], and could therefore be of importance in our model. Furthermore, combining recombination with co-infection of the simulated hosts would result in a model that allows one to study the emergence of recombinant HIV-1 types, which we intend to do in future work.

Probably even more important than the aforementioned limitations, is that it has proven difficult to construct a within-host ODE model of HIV-1 with both a realistic range of SPVL in the population, and a high heritability of this SPVL. Heritable variation in SPVL is important for our model because it facilitates efficient selection. The many-genes method [cf. 65] gives reasonable results, but our SPVL range is still 2 orders of magnitude narrower than observed ranges. In part, this is because various biological mechanisms are not incorporated in our model. As an example, including HIV-1 tropism and hosts with the CCR5-Δ32 allele could account for some of the lower SPVLs, thereby widening the SPVL distribution. Alternatively, the “shifting-mosaic metapopulation” model [54] could be used to get better results, but incorporating mutation and competition in such a model, while keeping the population-level model feasible to simulate will be challenging.

An important obstacle to curing HIV-1 is the latent reservoir: a population of infected, but quiescent CD4^+^ T cells that are invisible to the immune system and insensitive to antiretroviral drugs. This reservoir could be key to resolving short-sighted evolution with the optimization of ℛ_0_ [56]. This idea has recently been successfully explored by Doekes *et al*. [24] with the aid of a nested multi-strain model. Their model gave the best results when ancestral strains were preferentially transmitted, which in turn can be understood in light of a selection bias at the transmission bottleneck [16], and would explain the discrepancy between within-host and between-host evolutionary rates of HIV-1 [1, 55].

Finally, in spite of our strive for increased realism, we have used a relation between virus load and the duration of the infection that is still phenomenological. In a truly mechanistic model, the risk of AIDS would increase as the CD4^+^ T-cell count gets close to 200 cells per µl blood. Unfortunately, it is still not completely known how an untreated HIV-1 infection leads to progressive loss of CD4^+^ T cells. Interestingly, the rate at which the CD4^+^ T-cell count declines may be a heritable trait of the virus [7], in a manner that is independent from the heritability of the SPVL. This may lead to evolution of tolerance [74] and an increase of viremic non-progressors. We anticipate that such scenarios can be explored with a future version of the current model.

Concluding, our multi-level model allows us to perform large-scale *in silico* experiments that would be impossible to do otherwise. Our experiments show some interesting similarities with real-world observations, such as the emergence and impact of pre-adaptation, and the heritability of the SPVL. Naturally, many biological aspects are still missing—despite our aim for fidelity—some of which we have discussed above. We envisage that incorporating additional biologically relevant features in our current model could help answer other questions, including quantifying the impact of recombination, co-infection, and vaccination on viral evolution.

## Methods

### The individual-level model

#### The differential equations

At the core of the individual-level model, we have a system of ordinary differential equations (ODEs, see Equations (1)). The variables in this system of ODEs represent population sizes of quiescent CD4^+^ T-cells (Q), target cells (T), infected cells (I), CD8^+^ T cells, i.e. the effector cells (E), and free virus particles (V). When a target cell is infected, it first enters an eclipse phase (I_1_), before it starts to produce virions (phase I_2_). A graphical representation of the system is given in Figure 7A.

**Figure 7:**
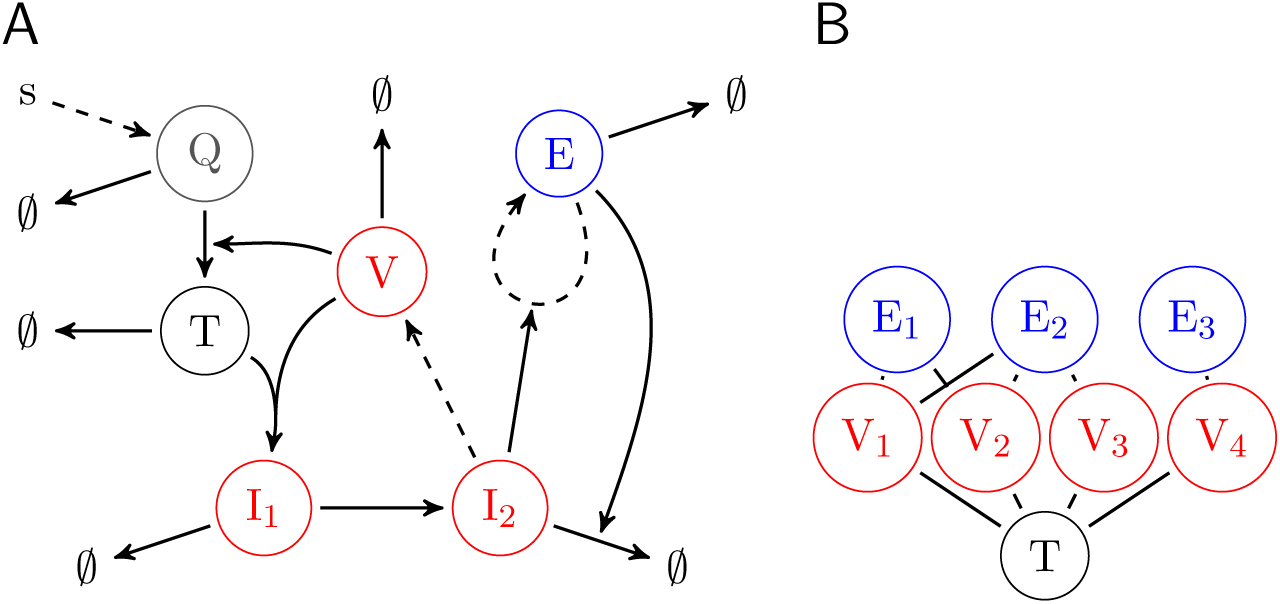
Graphical representation of the ODEs. **A**. In this example, only 1 viral strain and 1 immune response is present. Quiescent CD4^+^ T cells (Q) are produced from a source (s), and become activated target cells (T) at a rate that is dependent on the number of free virions (V). Target cells can be infected by virions, thereby producing eclipse-phase infected cells (I_1_). Eclipse-phase infected cells become virion producing cells (I_2_), that enhance the proliferation of effector cells (E). All populations are subject to *per-capita* death or clearance rates (∅), and the death-rate of virion-producing cells is enhanced by effector cells through killing. Production is indicated by a dashed arrow, while thick arrows signify a transition of catalysis. **B**. Virus strains and immune responses form a bipartite graph. In this example, 4 virus strains are controlled by 3 immune responses. Virus strain 4 is escaped from response 2, but the mutated epitope is recognized by response 3.

**Figure 8:**
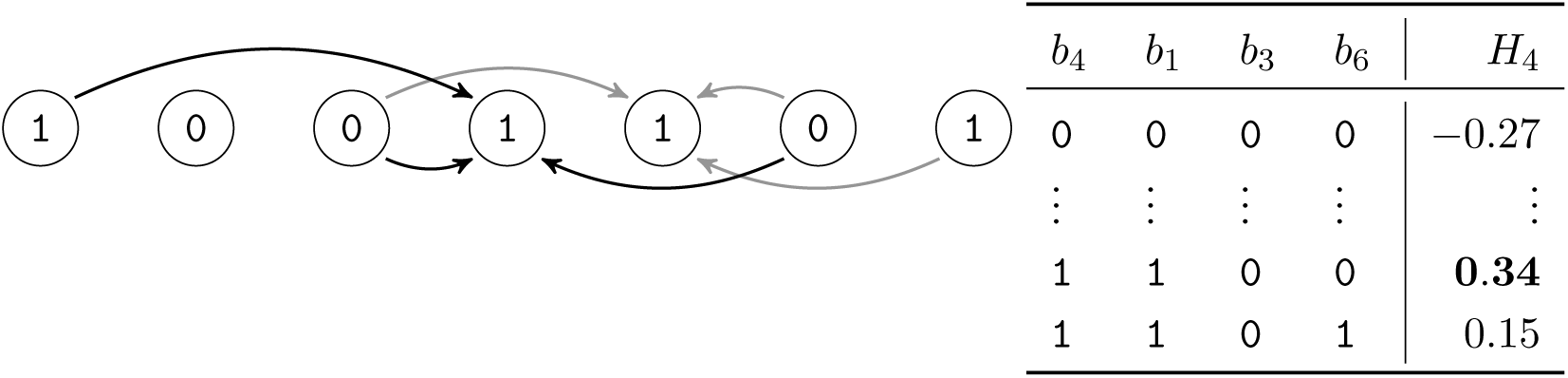
Computing a fitness parameter *w* from a bit string. In this example, we have *N* = 7 and *K* = 4. The black and gray arrows indicate the neighbors of locus 4 and 5, respectively. Locus 4 has neighbors 1, 3 and 6 (i.e. *U*_4_ = 4, 1, 3, 6). Given the alleles at locus 4 and its neighbors, we can find that the free energy *H*_4_ for this locus equals 0.34, from a pre-determined table. The total free energy is then given by 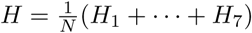, and the fitness parameter equals 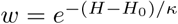.

The collection of all CTL clonotypes and virus strains form a bipartite graph *G* = (𝒱 ∪ ℛ, *ε*), with vertices corresponding to immune responses *r* ∈ ℛ and virus strains *v* ∈ 𝒱, and we draw an edge (*r, v*) ∈ *ε* whenever the response *r* recognizes an epitope of *v* (see Figure 7B). For a vertex *x*, we write 𝒩 (*x*) for the set of neighbors of *x*.

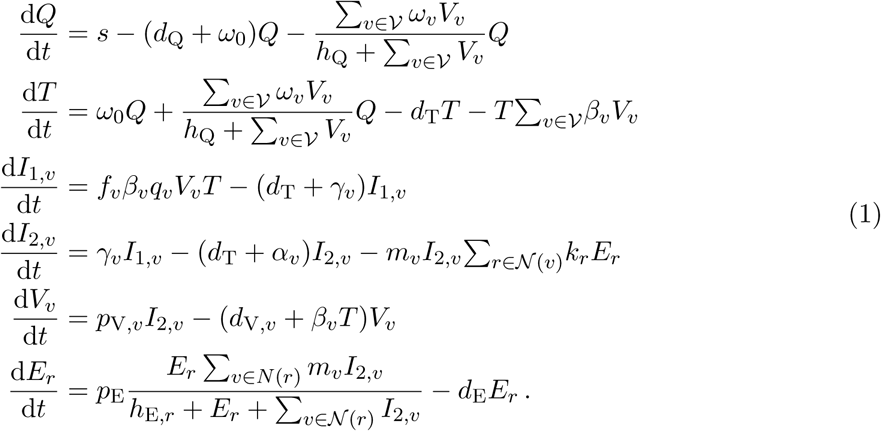

Because the dynamics of free virus particles is fast compared to the dynamics of infected cells, we make the quasi-steady-state (QSS) assumption 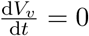, so that

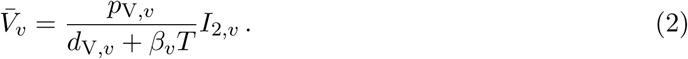

#### Evolution of the interaction graph

In addition to active responses and the number of infected cells, we also keep track of latent immune responses and all the single-locus mutants of the viruses present in the system. We think of these as stochastic elements of the system, as they are not described by deterministic ODEs.

#### Addition of novel viruses and immune responses to the interaction graph

In order to determine when a latent response or a mutant virus should be added to the system, we follow a scheme based on the Sellke algorithm [76]. This scheme consists of the following steps.

At time *t*_0_, we first sample a threshold *τ* ∼ Exp(1). We then add a differential equation for a load *L* to the system, and make the stochastic element deterministic when *L* ≥ *τ*. The initial value problem for *L* equals

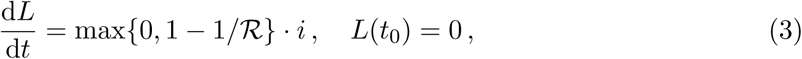

where ℛ is the reproduction number, and *i* is the incidence of either cells infected with the mutant, or activated CD8^+^ T cells. The factor max{0, 1 − 1*/*ℛ} in Equation (3) equals the so-called “major outbreak” probability of the stochastic population; it is the probability that a birth-death process does not reach the state of 0 individuals. Hence, we correct for the possibility that the newly generated cell clone can go extinct before its population size is large enough to be described by an ODE. Notice that this method is still somewhat phenemenological, as the true emergence dynamics of, e.g., escape mutants likely is much more complex [37].

The incidence of a mutant virus *v*′ is determined by the incidence of its parents, i.e. all virus strains that differ by only one locus from the mutant.

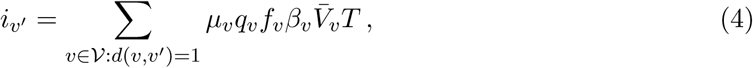

where *d* denotes the Hamming distance between viruses. The reproduction number of the mutant *v*′ equals

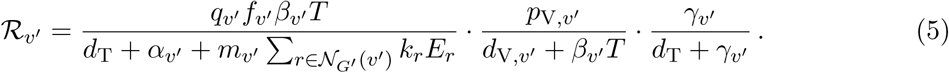

Here, 𝒩_*G*′_ (*v*′) denotes the set of responses in *G* that recognize an epitope of *v*′. Notice that the second factor in Equation (5) equals the number of virus particles per virion-producing cell. When the virus population becomes more diverse, we have to keep track of a large number of stochastic strains. For instance, the population gets extremely diverse during the acute phase, when the target-cell population is not yet depleted. To keep the simulations computationally feasible, we only follow stochastic strains that are mutants of deterministic viruses whose population sizes have at some point crossed a fixed threshold *τ*_mut_. In the simulations, we take *τ*_mut_ = 50 cells mL^−1^ (Figure 1).

A similar procedure is employed to determine when to add a naive or memory response to the system. The incidence of such a latent immune response *r*′ is constant, and differs only for naive and memory responses:

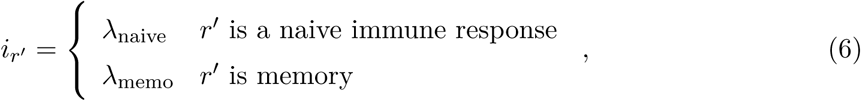

where we choose *λ*_memo_ ≫ *λ*_naive_ to express the fact that memory T cells are activated more rapidly than naive T cells. The reproduction number of a latent response *r*′ equals

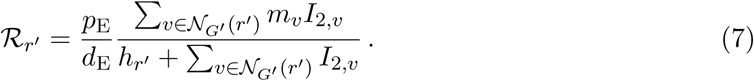

Here, 𝒩_*G*′_ (*r*′) denotes the set of viruses in *G* that contain the epitope that *r*′ recognizes.

### Deletion of extinct viruses and immune responses from the interaction graph

The ODEs give a deterministic description of the system, that is only a good approximation when the populations are large. When the size of a clone is small, we have to consider the risk of extinction. Whenever a clone size *N* (i.e. virus *I*_*v*_ or response *E*_*r*_) is below a threshold *τ*_end_, and its reproduction number ℛ is smaller than 1, the clone is considered endangered. When we find an endangered clone of size *N*_0_ ≈ *τ*_end_, we make a prediction about how long it takes for the clone to become extinct—*caeteris paribus*. Given that the clone has per-capita birth rate *b* and death rate *d*, a stochastic description of *N* ∈ ℤ_≥0_ is given by the following continuous-time Markov chain

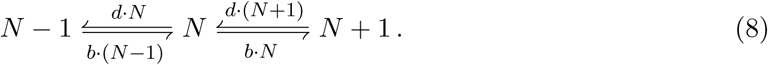

The goal is to compute the distribution of the “hitting time” *θ* of the state 0. This can be done by considering the probability generating function (PGF) of *N* [see e.g. 43, for more details]

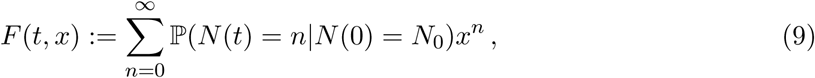

which satisfies the following PDE with boundary condition

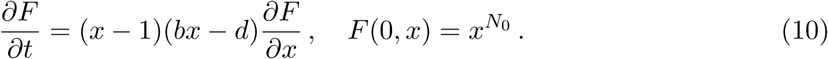

Using the method of characteristics, one finds that the distribution of the hitting time 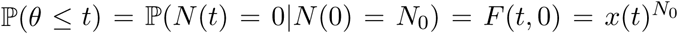, where *x* is the solution to the initial-value problem

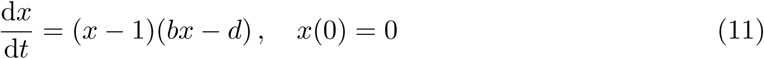

Therefore, we find that

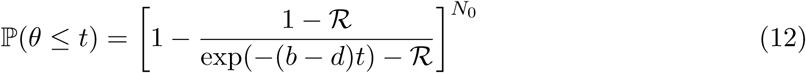

Notice that the deterministic description of *N* is approximately given by 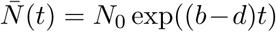 and hence we can write

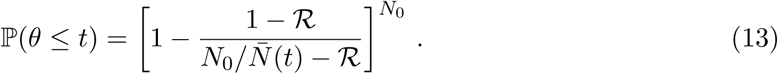

In order to decide when to remove a clone from the system of ODEs, we observe that for a random time *t* ∼ *θ*, the statistic ℙ(*θ* ≤ *t*|*t*) is uniformly distributed. Hence, we first sample a random deviate *u* ∼ Uniform(0, 1), and then solve Equation (13) for 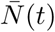. We remove the clone whenever its deterministic size drops below the extinction threshold:

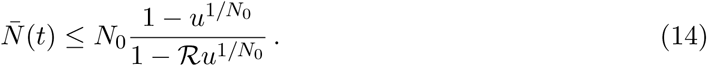

When a clone is endangered, we determine the extinction threshold only once. However, when the environment changes afterwards, the corresponding hitting time would no longer be valid. Since the threshold does not consider a particular time, but involves a population size, this issue is partially resolved. Consider, for example, the situation where a virus strain is endangered because of target-cell limitation. Before the clone reaches the extinction threshold, the target cell population might have recovered, such that the actual growth rate of the endangered strain is larger than the growth rate at the time when the extinction threshold was determined. This means that extinction will occur later than anticipated. This, however, is qualitatively consistent with the stochastic description. Whenever an endangered clone grows above the threshold *τ*_end_, we consider it rescued, and if thereafter it falls below the threshold *τ*_end_ again, we randomly draw a new extinction threshold. In the simulations, we take *τ*_end_ = 10 cells mL^−1^ (Figure 1).

We use this scheme to remove both active immune responses and viral strains from the system of ODEs. Notice, however, that a viral strain consists of a multi-type population, i.e. I_1_, I_1_ and V, where we make a QSS assumption for *V*. In order to use Formula (14) we simply use the sum of the infected cells in eclipse and producing stages: *N* = *I*_0_ + *I*_1_.

### Host and virus contributions to parameter variation

The parameters of the ODEs depend on the virus, immune responses, and the remaining genetic make-up of the host. A parameter *x*_*v*_ that is determined by both host and virus is calculated as follows. For a host we draw *x*_host_ from a Log-Normal distribution with mean *x*_default_ and a standard deviation of 10% [65]. The virus at node *v* has a gene that determines a fitness parameter *w*_*x*_ via the *NK* map (see below). For the virus we compute a parameter *x*_virus_ = *x*_default_ · *w*_*x*_, whenever a higher value of the parameter corresponds with a higher fitness (e.g. *x* = *β* or *p*_V_), and *x*_virus_ = *x*_default_*/w*_*x*_ when a higher value of the parameter results in a lower viral fitness (e.g. *x* = *α* or *d*_V_). The parameter *x*_*v*_ that is used in the system of ODEs is the geometric mean of *x*_virus_ and *x*_host_, i.e., 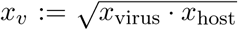. We use the geometric mean instead of the arithmetic mean because otherwise the value of the parameter would be limited from below by the host. For example, when a non-viable mutation would result in *β*_virus_ ≈ 0, then we require *β*_*v*_ ≈ 0, which is true for the geometric, but not the arithmetic mean.

The parameters of the ODEs that are co-determined by the virus are derived using the *NK* model that provides a genotype-to-phenotype map [44]. In this model, a genome is modeled using a string of *N* bits *b*_*ℓ*_. For each locus *ℓ* of the genome, a local neighborhood *U*_*ℓ*_ of *K* loci is chosen, and the alleles *b*_*k*_ at the loci *k* ∈ *U*_*ℓ*_ together determine the energy 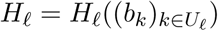 associated with locus *ℓ*. The total energy *H* of the genome 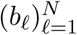 is defined as the average of the local energies: 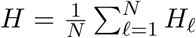. For each local neighborhood *U*_*ℓ*_ we have to determine 2^*K*^ energies, one for each genetic background 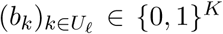. These values are generated at random at the initialization step of the simulation, and are 𝒩(0, 1) distributed.

The neighborhoods *U*_*ℓ*_ are chosen as follows. First, we choose *K*_*ℓ*_ ∼ Uniform(1, 2, …, *K*_max_) with *K*_max_ = 10 and we always let *ℓ* ∈ *U*_*ℓ*_. Then we choose *K*_*ℓ*_ − 1 unique neighbors *k* = *ℓ* + *X* where *X* ∼ Skellam(*L, L*), and *L* = 3*K*_*ℓ*_. We have a separate gene for each parameter, which is a consecutive sub-string of the genome. All genes have the same length *N*, and only loci in the gene contribute to the parameter-specific energy. The neighborhood of a gene’s locus can contain loci of nearby genes on the genome, which facilitates epistatic interactions between genes. Such interactions have for instance been observed between Gag and Protease [28]. The energy *H* is transformed into to a positive, dimensionless number *w* = exp(−(*H* − *H*_0_)*/κ*), where *κ* plays the role of the temperature and can be used to tune the ruggedness of the fitness landscape. By trial and error, we found that *κ* = 0.1 results in reasonable fitness costs.

The parameter *H*_0_ is used to determine for which energies *H* the virus-effect on the parameter is neutral (i.e. *w* ≈ 1). For large *N*, we have 𝔼[*H*] = 0, for random genomes, but for evolved genomes, *H* can be substantially larger than 0. We therefore start with *H*_0_ = 0, and sample a large number of random genomes. These genomes are then evolved using a hill-climbing algorithm, resulting in a sample of evolved energies, and *H*_0_ is henceforth defined as the median of these energies.

### Immune responses, epitopes, and HLA alleles

The same string of bits that is used as a model of a genome, is used to encode the epitopes, i.e. the targets of the immune responses. Each immune response *r* targets an epitope at a fixed locus *ℓ*_*r*_ in the genome. The epitope consists of 3 consecutive bits, where the first bit serves as an anchor position, i.e. this bit models the amino acids that are largely determining binding with the HLA molecule. The remaining bits determine TCR binding. Hence, each epitope has 2^2^ = 4 variants, and at each locus, there are potentially 2 epitopes, one for each anchor bit. In reality, the number of epitopes is much larger, but TCRs are also known to cross-react, i.e. are able to induce a T-cell response to multiple variants of the same epitope, or even multiple epitopes. Additionally, the motifs corresponding to HLA alleles can allow for multiple amino-acids at the anchor positions, which also reduces the effective variability of epitopes.

Immune responses can be shared between individuals, and determine 2 parameters of the system of ODEs. The parameter *k*_*r*_ determines the killing rate of immune response *r*, and the Michaelis-Menten parameter *h*_E,*r*_ determines the amount of antigen that is required for response *r* to expand. Since we expect *k*_*r*_ and *h*_E,*r*_ to be correlated, we first choose an immuno-dominance parameter 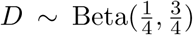, such that sub-dominant responses are more common, and then choose weighted geometric averages 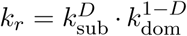 and 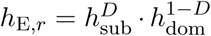, after which we add 10% Log-Normal noise.

A HLA allele is modeled as a set of responses against a fixed set of epitopes. On average, we take 10 epitopes per HLA allele (with a 25% standard deviation), and hence 40 = 2^2^ × 10 responses against all variants of these epitopes. Each host has 6 = 3 × 2 HLA alleles (3 HLA class I loci on each of the 2 chromosomes). Each HLA locus has 30 alleles with a fixed allele frequency distribution, which is sampled from the Dirichlet(1, …, 1) distribution, giving rise to both common and rare alleles.

### The population-level model

#### A dynamic contact network

We simulate a human population where HIV-1 spreads through heterosexual transmission. In our design of the contact network, we aimed for the simplest model that at least has some features that are inherent to real human populations: (i) Individuals have sexual contacts with a limited number of individuals within a time window, but this number differs between individuals, resulting in different risk groups in the population. (ii) Contact formation is assortative as depends typically on age difference, and similarity of promiscuous behavior. (iii) Contact networks display a certain amount of clustering.

The simulated population consists of individuals, or agents, and contacts between these agents (see Figure S1). The duration of a contact is finite and Log-Normal-distributed, with a mean and standard deviation of 1y. A subset of the contacts can be used as a transmission route (sexual contacts). Agents have an intrinsic contact-formation rate *λ*_cf_ (which is Log-Normal-distributed, mean = 2y^−1^, sd = 1y^−1^), and attempt to form new contacts according to a Poisson process with intensity *λ*_cf_. The maximum number of sexual contacts *K*_tmr_, differs between individuals and determines their promiscuity. We let *K*_tmr_ ∼ Pos-Poisson(*λ*_cf_), where Pos-Poisson is a zero-truncated Poisson distribution, such that individuals with a higher contact formation rate also tend to have more concurrent sexual contacts.

When an agent *A* requires a new contact, it first searches for a new partner in their local neighborhood, which is defined as the set of agents *B* such that *A* and *B* share a partner [cf. 62]. The probability of contact formation depends on age difference Δ*a*, and the difference in the level of promiscuity Δ*K*_tmr_:

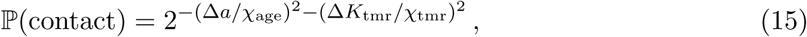

where *χ*_age_ and *χ*_tmr_ are the characteristic age difference and promiscuity difference at which contact the contact formation probability ℙ(contact) is halved. When no compatible contact was found in the local neighborhood, a new partner is sought in the total population, again giving precedence to contacts with a higher ℙ(contact).

Each agent always tries to maximize its number of sexual contacts. In order to accomplish this, we walk through the population in a random order. For each agent, we test for each contact (again in a random order) if it can become a sexual contact. This is allowed if: (i) the partners have opposite gender, (ii) both partners are older than the minimum age, (iii) both partners have fewer sexual contacts that their limit *K*_tmr_.

#### Demographic turn-over

Susceptible individuals enter the population at a constant rate *λ*_birth_. The lifespans of susceptible individuals that are entering the population are sampled from a realistic parametric distribution with hazard *x* 1↦ *e*^0.1*x*−10.5^ +*e*^−0.4*x*−8^ [17]. The initial population is seeded with ages sampled from the distribution that corresponds to the above hazard, assuming a demographic equilibrium. The hazard is also used to compute the birth rate *λ*_birth_, such that a virgin population has the desired size.

#### Transmission and HIV-1 induced mortality

Transmission of the virus is dependent on the viral load. The transmission rate *β* of an infected individual with viral load *V* is given by the Hill function

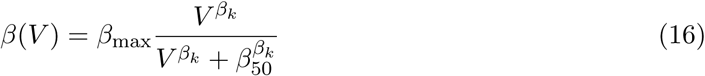

where *β*_max_ is the maximal transmission rate, the parameter *β*_50_ is the virus load for which the transmission rate is 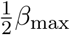, and *β*_*k*_ is a shape parameter. Notice that we use the symbol *β* for both the individual-level model infection rate as the population-level model transmission rate. The values for these parameters were estimated using a cohort of transmission couples [29, and see Table 2]. For determining the moment that a susceptible individual gets infected, we again use the Sellke construction [76]. A susceptible individual *i* accumulates the sum of all transmission rates from all its sexual contacts *j*, until the cumulative transmission rate exceeds a pre-determined threshold *τ*_infect_ ∼ Exp(1). More precisely, we define a infection load *L* via

**Table 2:** Parameters of the population-level model. Notes: The parameters (1) are taken from Fraser *et al*. [29] and (3) Shirreff *et al*. [77]. (2) The sexual contact network are chosen by trial and error, with the requirement that an HIV-1 outbreak is possible.

| symbol | description | value | unit | note |
| --- | --- | --- | --- | --- |
| $\beta_{\max}$ | maximum transmission rate | 0.317 | y<sup>-1</sup> | (1) |
| $\beta_{50}$ | VL at half maximum transmission rate | 13 938 | copies/ml | (1) |
| $\beta_k$ | shape parameter of the transmission rate | 1.02 | - | (1) |
| $D_{\max}$ | maximum expected infection length | 25.4 | y | (1) |
| $D_{50}$ | VL at half maximum expected infection length | 3058 | copies/ml | (1) |
| $D_k$ | shape parameter of the expected infection length | 0.41 | - | (1) |
| $\rho$ | shape parameter of the distribution of the infection length | 3.46 | - | (1) |
| $\lambda_{\text{cf}}$ | contact formation rate | 2 | y<sup>-1</sup> | (2) |
| $\chi_{\text{age}}$ | age difference at half maximum contact formation probability | 12 | y | (2) |
| $\chi_{\text{tmr}}$ | promiscuity difference at half maximum contact formation probability | 1 | partners | (2) |
| $\sigma_M$ | mutational variance | 0.12 | log<sub>10</sub> copies/ml | (3) |

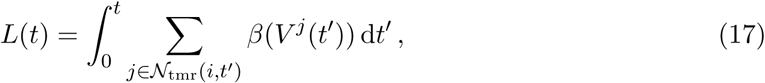

and infect the susceptible host at time *t*_infect_ := inf{*t* : *L*(*t*) ≥ *τ*_infect_}. Here, 𝒩_tmr_(*i, t*) is the set of all sexual contacts of *i* at time *t*. This process is approximated using a *τ* -leaping algorithm with a time step of 1 week. The transmitter is chosen from all infectious partners with probability proportional to the partners’ transmission rates *β*(*V* ^*j*^). From the transmitter, in turn, a virus *v* is sampled with probability proportional to the clone-specific virus load 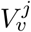.

After an acute and asymptomatic infection, an infected agent progresses to AIDS and dies. The duration of the asymptomatic infection depends on the virus load. For a constant virus load *V*, the average duration of the chronic phase equals

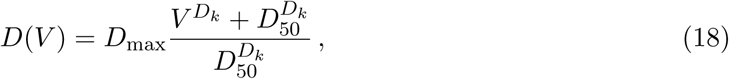

Conditioned on the constant virus load, observed lengths of chronic phases show significant variation and can be modeled with a Gamma distribution with mean *D*(*V*) and shape parameter *ρ*. These parameters were estimated from a cohort of untreated individuals [29, and see Table 2]. Since the virus load on an infected individual is not constant, we interpret the reciprocal duration *D*(*V* (*t*))^−1^ as a hazard of progressing to AIDS, and remove an infected individual from the population whenever its cumulative hazard 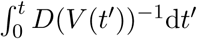 exceeds a pre-determined threshold *τ*_death_ ∼ Gamma(*ρ, ρ*^−1^). Importantly, when individuals are “cloned” to determine the effect of immunological pre-adaptations, the threshold *τ*_death_ is inherited by the clone. Notice that an HIV-1 induced death event can be censored by the natural death event of the infected individual, and vice versa. However, all individual-level statistics (such as the SPVL) are always calculated taking the HIV-1 induced death event as the end of the simulation.

To compare our results with the mutation-at-transmission model [77], a virus is given a genome-independent trait *g* that modulates the log_10_ viral load. In order to determine AIDS hazard and infectiousness, the viral load log_10_(*V*) is replaced by log_10_(*V*) + *g*. The viral load used for the individual-level model remains unchanged. The trait *g* is allowed to mutate when the virus is transmitted using mutational variance *σ*_*M*_ = 0.12 [77], i.e. the transmitted virus samples a 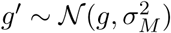.

### Associations between HLA alleles and viral polymorphisms

In order to discover associations between HLA alleles and viral polymorphisms, we take a similar approach as, e.g., Carlson *et al*. [15]. *Let b*_*ℓ*_ be the allele at locus *ℓ*, and let *b*_*ℓ*,WT_ be the wild-type allele. We want to test if a mutation at locus *ℓ* happens preferentially in the context of an HLA allele *i*. Let *X*_*i*_ ∈ {0, 1} represent the absence or presence of HLA allele *i*. The probability *P*_*ℓ*_ that *b*_*ℓ*_ ≠ *b*_*ℓ*,WT_ is modeled as

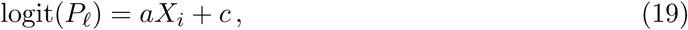

where logit(*x*) := log(*x/*(1 − *x*)) and parameters *a* and *c* are estimated by maximizing the likelihood. The parameter *a* equals the log-odds that the mutation occurs due to the presence of HLA allele *i*. The mutation can also be observed because it is present in the transmitted virus. In order to correct for this bias, we define a variable *T*_*ℓ*_ ∈ {−1, 1} that equals 1 if the mutation is transmitted. The corrected model equals

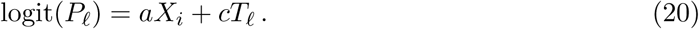

For each locus *ℓ* and HLA allele *i*, we compute the *p*-value under the null hypothesis that *a* = 0 using the likelihood-ratio test. Pairs (*ℓ, i*) for which any of the elements of the contingency table is less than 5 individuals are ignored. In order to correct for multiple testing, the *p*-values are transformed into *q*-values using the procedure described by Storey and Tibshirani [79]. An association is called significant if *p <* 0.05 and *q <* 0.2.

### Implementation

The model is written in the `C++` programming language and the output of the simulations is analyzed with Python. The source code and scripts can be downloaded from www.github.com/chvandorp/mlev-hiv-model.

## Acknowledgments

This work has been funded by The Netherlands Organization for Scientific Research (NWO; grant numbers 645.000.002 and 823.02.014), and the National Institute for Public Health and the Environment (RIVM).

---

**Figure S1:**
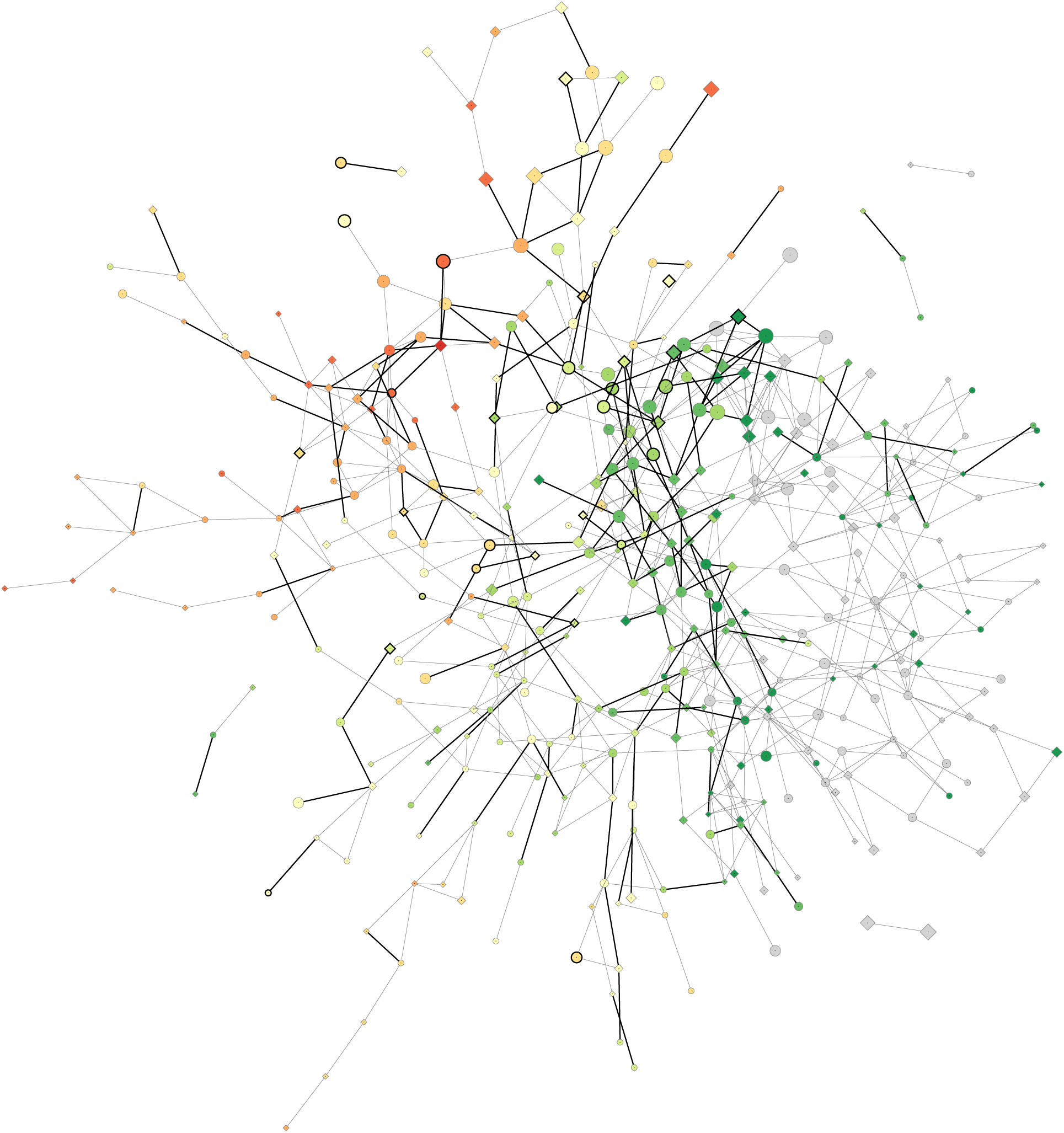
Snapshot of the dynamic contact network. Nodes represent the individuals and edges represent contacts between individuals. The heavy edges represent contacts that can be used as transmission routes (sexual contacts), while the light gray edges represent the auxiliary contacts (social contacts). Nodes with a thick outline represent infected agents, and the color of the node corresponds to the age of the individual. The gray nodes are individuals outside the population at risk (individuals of age ≤ 15y). The size of the node represents the promiscuity of the individual (the maximum amount of concurrent sexual contacts allowed). The shape of the node represents the gender (○ for males and ◊ for females).

**Figure S2:**
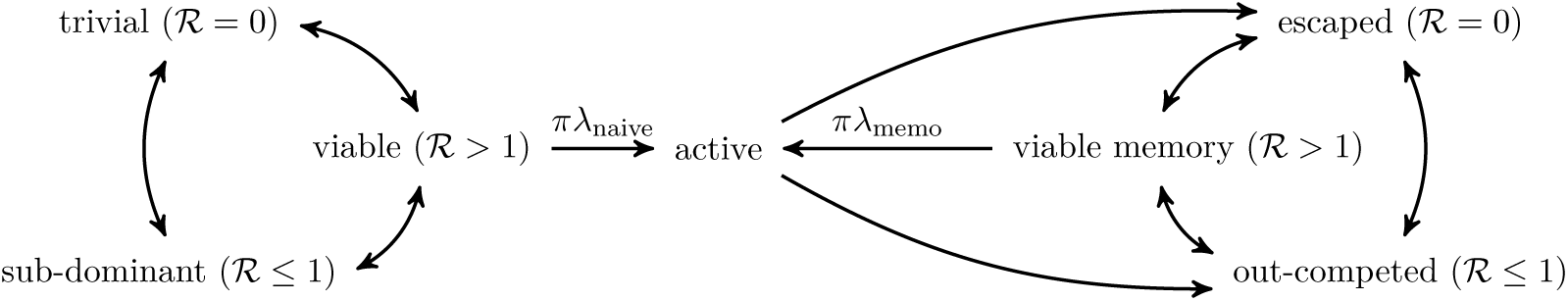
The states and transitions of immune responses. Naive responses can toggle between a trivial, sub-dominant and viable state, depending on their reproduction numbers ℛ. An immune response is in the trivial state when none of the strains contain its epitope. Viable responses are activated at rate *πλ*_naive_, where *π* = 1−1*/*ℛ. Active responses can either be escaped by a fixating escape mutant, or out-competed by a stronger response. In any case the response is now in a memory state. Out-competed responses can be escaped by chance, and memory response can become viable memory when the virus reverts the epitope, or when another strong responses is escaped.

**Figure S3:**
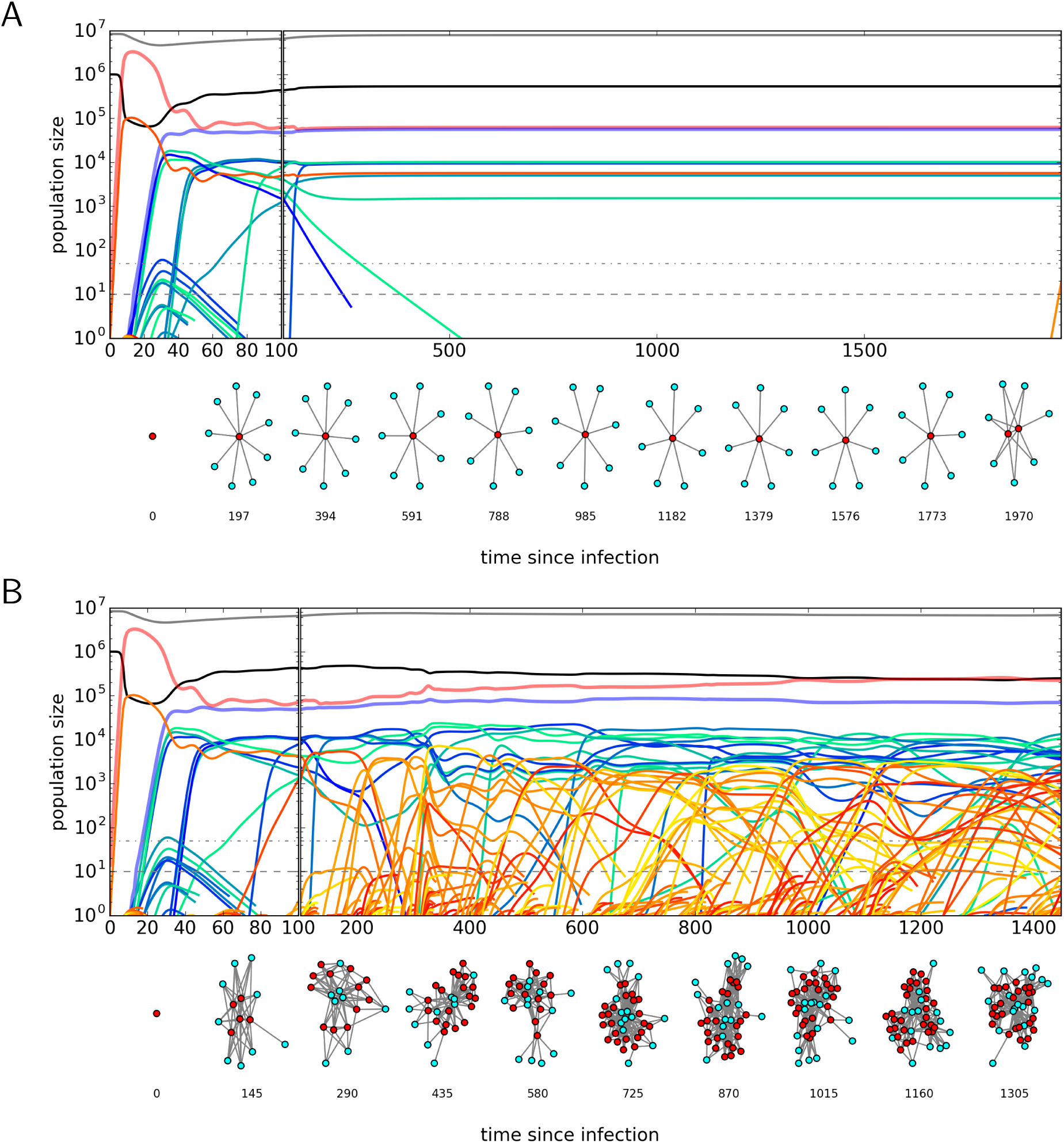
Individual-level simulations with low and high mutation rates. The parameters initial conditions are identical those used for Figure 1, with the exception of the mutation rate *µ*, which is tenfold lower in panel **A** (*µ* = 10^−7^), and tenfold higher in panel **B** (*µ* = 10^−5^).

**Figure S4:**
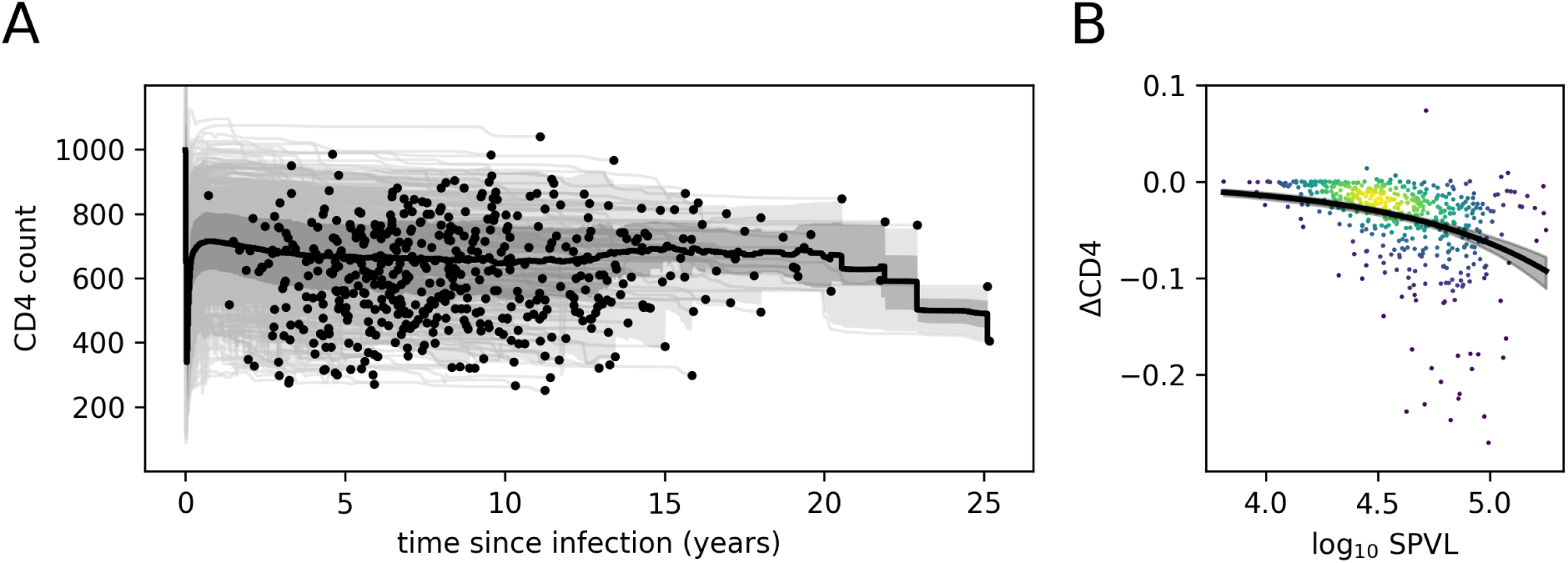
Relation between CD4^+^ T-cell decline and SPVL. The data is taken from a crosssectional sample of 500 patients, taken 200 years into the epidemic. **A**. A collection of individual-level time series of the CD4^+^ T-cell count (CD4 count: target cells per µl blood). The gray bands indicate the 2.5, 25, 75, and 97.5 percentiles, the thick line the average. The end of an infection is indicated with a black dot. **B**. The relation between log_10_ SPVL and the slope of the CD4^+^ T-cell count (ΔCD4: cells per µl per day). The hue indicates the density of the observations. An exponential curve is fitted through the data (by non-linear least squares). The gray bands indicate the 95% CI (calculated by bootstrapping the data).

**Figure S5:**
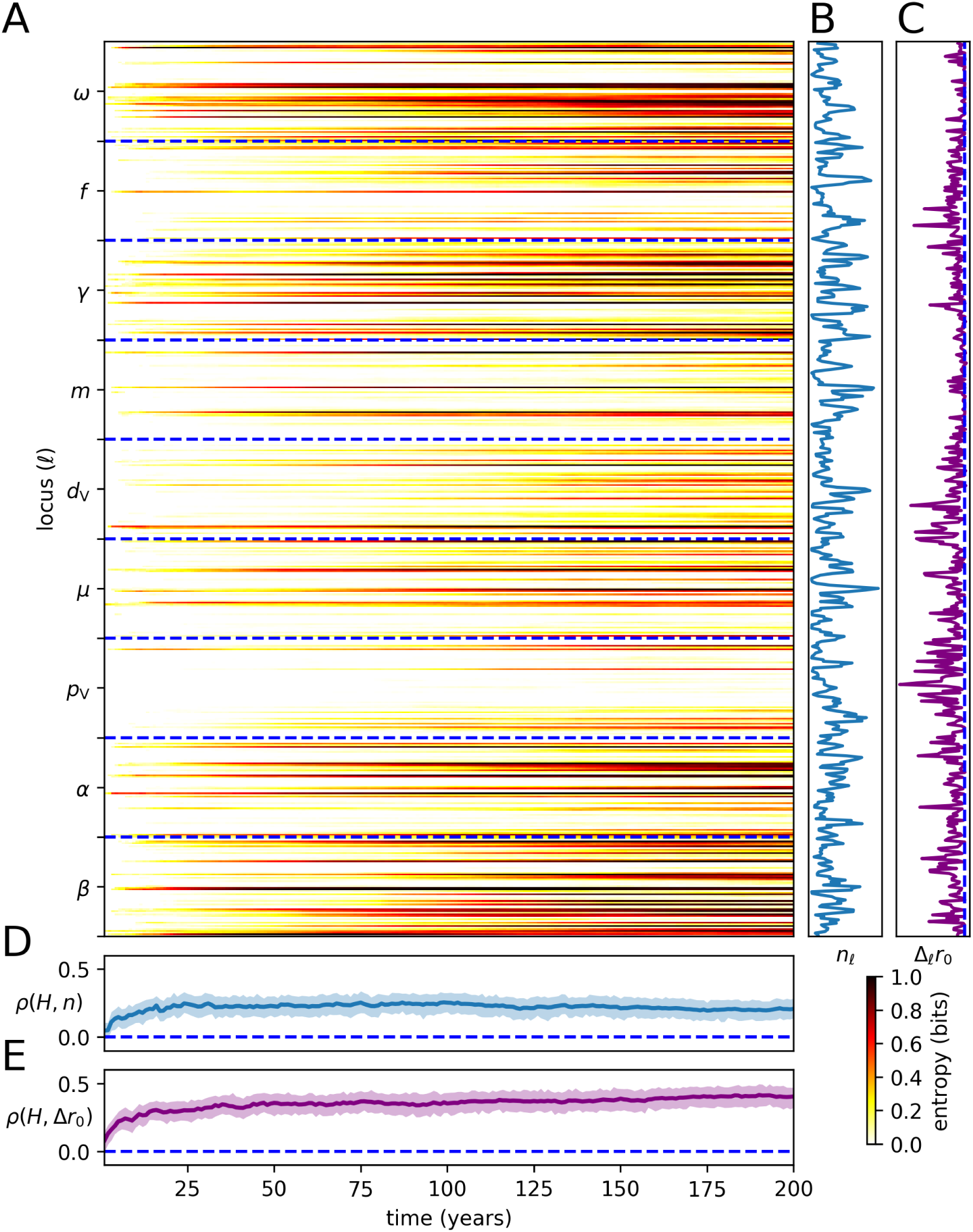
Genetic diversity of the virus per viral locus correlates with epitope density and fitness cost. **A**. For each simulated year and each locus *ℓ* of the genome, we compute the fraction of genomes *b* that have allele 1 at locus *ℓ*, taking all *I* infected individuals into account: 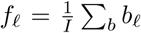. The corresponding entropy equals *H*_*ℓ*_ = *f*_*ℓ*_ log_2_(*f*_*ℓ*_). **B**. The number of responses *n*_*ℓ*_ targeting an epitope that contains the locus *ℓ* is weighted by both the fixed HLA frequency distribution, and the immuno-dominance parameter *D*. **C**. For each locus *ℓ*, we consider the replicative capacity of a transmitted virus and a virus with a point mutation at locus *ℓ*. The relative basic Malthusian fitness (Δ_*ℓ*_*r*_0_) is the difference between these vRCs. **D**. The weighted number of responses *n*_*ℓ*_ directed at locus *ℓ* is correlated with the entropy *H*_*ℓ*_ for each simulated year (using Spearman’s *ρ*). The colored band represents the 95% CI (calculated using bootstrapping). **E**. The relative basic Malthusian fitness (Δ*r*_0_) is correlated with the entropy.

